# SUMOylation of ABCD3 restricts bile acid synthesis and regulates metabolic homeostasis

**DOI:** 10.1101/2022.03.03.482848

**Authors:** Vanessa Goyon, Aurèle Besse-Patin, Rodolfo Zunino, Mai Nguyen, Étienne Coyaud, Jonathan M. Lee, Bich N. Nguyen, Brian Raught, Heidi M. McBride

## Abstract

Mitochondrial anchored protein ligase (MAPL) has been shown to function as both a SUMO and ubiquitin ligase with multiple roles in mitochondrial quality control, cell death pathways and inflammation. To examine the global function of MAPL we generated a knock-out mouse model and sought functional insight through unbiased BioID, transcriptomics and metabolic analysis. MAPL KO mice are lean and highly insulin sensitive, ultimately developing fully penetrant, spontaneous hepatocellular carcinoma after 18 months. BioID revealed the peroxisomal bile acid transporter ABCD3 as a primary MAPL interacting partner, which we show is SUMOylated in a MAPL-dependent manner. MAPL KO animals showed increased bile acid secretion in vivo and in isolated primary hepatocytes, along with robust compensatory changes in the expression of enzymes synthesizing and detoxifying bile acid. In addition, MAPL KO livers showed signs of ER stress and secreted high levels of Fgf21, the starvation hormone known to drive the reduction of white fat stores and promote insulin sensitivity. Lastly, during aging all MAPL KO mice developed hepatocellular carcinomas. These data reveal a major function for MAPL in the regulation of bile acid synthesis leading to profound changes in whole body metabolism and the ultimate generation of liver cancer when MAPL is lost.

## Introduction

Mitochondria are an essential signaling platform that contributes to cell fate decisions, from cell cycle transitions to stem cell differentiation, T-cell activation, pathogen invasion, starvation and more (Nunnari and Suomalainen, 2012). An underlying reason for the integration of mitochondria within cellular signaling pathways is to signal the transcriptional and post-transcriptional changes required to alter fuel consumption and/or metabolite generation that drive cellular transitions. In this way signaling at mitochondria contributes to the global rewiring of cellular metabolism in response to a variety of extracellular stimuli and intracellular cues. It is becoming apparent that, akin to other signaling pathways, post-translational modifications including phosphorylation, ubiquitination and SUMOylation play a central role in the assembly and regulation of mitochondrial signaling complexes (Escobar-Henriques and Langer, 2014; He et al., 2020; Tait and Green, 2012; Tan and Finkel, 2020). The mechanistic details responsible for these modifications in mitochondrial signaling are however still largely unknown.

MAPL, a mitochondrial and peroxisomal anchored protein ligase (also called MUL1/GIDE/HADES/MULAN) (Jung et al., 2011; Li et al., 2015, 2008; Zhang et al., 2008) is a mitochondrial outer membrane protein carrying two transmembrane domains, a ~40kDa intermembrane space loop and C-terminal cytosolic RING finger with both SUMO and ubiquitination activities. It is also targeted to peroxisomes in mitochondrial vesicles and is part of the shared peroxisome/mitochondrial proteome. MAPL has been shown to modulate diverse cellular events including mitochondrial division, mitophagy, inflammation and cell death (Ambivero et al., 2014; Barry et al., 2018; Prudent et al., 2015; Rojansky et al., 2016; Yun et al., 2014). A common feature of MAPL induced SUMOylation of substrates is to stabilize complex formation or assembly, making it a prime candidate regulator of signaling platforms on mitochondria and peroxisomes (Prudent et al., 2015).

To better understand the primary function of MAPL we have explored the proximity interactors using unbiased BioID approaches (Roux, 2013; Roux et al., 2012). In addition to the core mitochondrial and peroxisomal fission machinery, the BioID identified peroxisomal ABCD3 as an interacting partner and SUMOylation substrate of MAPL. ABCD3 was shown to transport the late stage precursors, the C27-bile acid intermediates 3α,7α-dihydroxycholestanoic acid (DHCA), 3α,7α,12α-trihydroxycholestanoic acid (THCA), and dicarboxylic fatty acids from cytosol into peroxisomes (Ferdinandusse et al., 2015; Ranea-Robles et al., 2021). With the generation of a MAPL knock-out mouse model, we uncovered a critical role for this SUMO E3 ligase in restricting the activity of ABCD3, highlighting new links to whole body metabolism. Further analysis of the MAPL deficient mice revealed increased hepatocyte proliferation, resistance to programmed cell death, and the development of hepatocellular carcinoma in aging mice. These data provide new insights into the post-translational regulation of bile acid metabolism within the liver, and the central role for peroxisomal SUMOylation in metabolic homeostasis.

## Results

### ABCD3/PMP70 is a MAPL substrate

To identify MAPL interacting proteins we performed an unbiased proximity-dependent biotin identification (BioID) in HEK293 cells (Roux, 2013; Roux et al., 2012). As expected, BioID confirmed a robust interaction between MAPL and the fission GTPase DRP1 **(Fig 1A)** (Braschi et al., 2009; Neuspiel et al., 2008; Prudent et al., 2015). Indeed, essentially all of the mitochondrial fission machinery identified to date was identified in this analysis, including: the regulator of DRP1 recruitment AKAP1, the cAMP-dependent protein kinase type II-alpha regulatory subunit PRKAR2A, the DRP1 receptor MFF, the inverted formin INF2 (Kraus and Ryan, 2017), USP30, a regulator of mitophagy and pexophagy (Bingol et al., 2014; Marcassa et al., 2018), and others. Also present was the antiviral signaling protein MAVS, an interaction we and others have characterized previously (Doiron et al., 2017; Jenkins et al., 2013). Potential ubiquitin substrates of MAPL such as MFN1, MFN2, AKT, HIF1α, or P53 were absent from the interactome at steady state.

**Figure 1:**
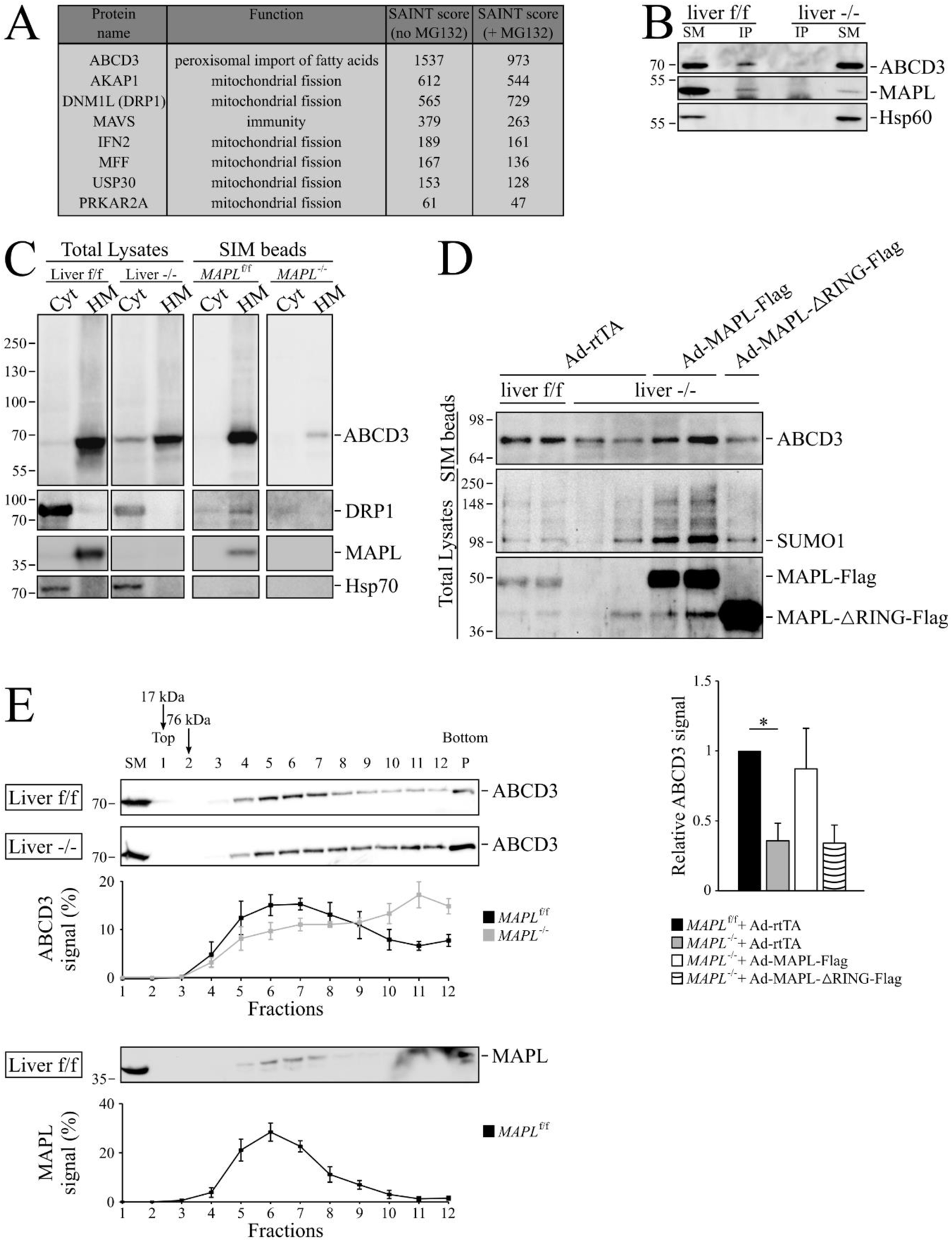
MAPL SUMOylates ABCD3 and modulates complex assembly. **A.** HEK293T-REX (tetracyline-regulatable expression) cells stably expressing an inducible Tet-ON fusion construct MAPL-Flag-BirA or Flag-BirA were induced for 24 hours in the presence of biotin, and biotinylated proteins were isolated with streptavidin beads for identification by mass spectrometry. The top hits by peptide counts are shown. **B.** Starting materials (SM) and MAPL immunoprecipitated (IP) fractions obtained from liver crude extracts were probed for ABCD3, MAPL and Hsp60. **C.** Cytosol and heavy membrane fractions isolated from MAPL^f/f^ and MAPL^−/−^ livers were solubilized and incubated with SIM beads, and elution fractions probed for ABCD3, DRP1, Hsp70 and MAPL (Cyt=cytosolic fraction, HM=heavy membrane fraction). **D.** Heavy membrane fractions isolated from livers of MAPL^f/f^ and MAPL^−/−^ animals tail-vein injected with adenovirus expressing MAPL-Flag, MAPL-ΔRING-Flag or empty virus (Ad-rtTA) were solubilized and incubated with SIM beads. Elution fractions were probed for ABCD3, SUMO1 and MAPL (top panel). Quantification from 3 independent experiments of ABCD3 signals of the heavy membrane SIM-beads elution fraction from livers isolated from rescued mice (lower panel). **E.** 250 µg of solubilized protein (from MAPL^f/f^ and MAPL^−/−^ livers) were separated on a 10-50% (w/v) sucrose gradient. 12 different fractions, as well as the resuspended pellet (P) were analyzed by western blot. ABCD3 signals from MAPL^f/f^ and MAPL^−/−^ livers in the different fractions were quantified and plotted as percentage of the total signal, from 3 biological replicates. * *P* < 0.05 in a one-way ANOVA

Many ubiquitin E3 ligases target their substrates for 26S proteasome-mediated degradation, perhaps explaining the absence of any expected ubiquitin substrates of MAPL. Therefore, we repeated the BioID experiments in the presence of MG132 over 24 hours to stabilize and accumulate any potential ubiquitinated MAPL-FlagBirA* substrates (Coyaud et al., 2015) (**Supplemental Table 1**). Notably, the number of peptides identified for the vast majority of MAPL interacting proteins remained unchanged in the presence of MG132, or were decreased, suggesting that MAPL does not target its primary binding partners for proteasomal degradation. A few additional proteins were detected at very low levels after 24 hours of MG132 treatment, with peptide counts ranging from 7 to24 (compared to 612 for AKAP1) including BAX, MIRO1, BNIP3 and MFN2.

Unexpectedly, the top ranked MAPL binding protein detected in this analysis was the peroxisomal bile acid transporter ABCD3/PMP70 **(Fig 1A)**. We confirmed this result using western blot analysis of biotinylated ABCD3 captured on streptavidin beads after incubation of MAPL-BirA expressing cells with biotin **(Supplemental Fig 1A)**. MAPL is delivered to peroxisomes through a vesicular transport pathway from mitochondria (Braschi et al., 2010; Neuspiel et al., 2008), where it plays a role in regulating peroxisomal fission (Mohanty et al., 2021). However, broader roles of MAPL in peroxisomes are unknown. Given that ABCD3 has a primary role in the generation of bile acids, we chose to validate the interaction and interrogate the potential functional consequences of their binding within the liver of our MAPL^−/−^ mouse line (Doiron et al., 2017). Briefly, a parental C57Bl/6J strain carrying floxed alleles at exon2 was crossed with a *CMV*-*Cre* carrying strain to excise exon 2 in all cells, including the germline. After backcrossing out the *Cre* gene, the resultant strain was a germline knock out for MAPL (Doiron et al., 2017). We used these mice to test any interaction between native proteins by immunoprecipitating endogenous MAPL from liver of control or MAPL*^−/−^* mice. While the anti-MAPL antibodies did not efficiently precipitate endogenous MAPL, we still observed a MAPL-dependent interaction with ABCD3 from liver tissue (**Fig 1B**). We next monitored the SUMOylation of ABCD3 within the liver of MAPL^−/−^ mice. For this we isolated cytosol (Cyt) and solubilized the heavy membrane (HM) fraction from MAPL^f/f^ and MAPL^−/−^ livers. These fractions were incubated with agarose beads conjugated to peptides encoding the consensus SUMO interacting motif (SIM) of the nuclear SUMO E3 ligases PIAS1-4 (Hecker et al., 2006). Indeed, we observed an ABCD3 immunoreactive band on the SIM beads of control livers, which was almost completely absent in livers from mice lacking MAPL (**Fig 1C, quantification Supplemental Fig 1B**). The molecular weight of ABCD3 upon the SIM beads was slightly shifted, suggesting a mono-SUMOylation event. As a positive control MAPL was also required for the SUMOylation of DRP1 (**Fig 1C**) (Braschi et al., 2009; Prudent et al., 2015).

To further confirm whether MAPL was responsible for the SUMOylation of ABCD3 we generated adenovirus expressing MAPL-Flag, or a deletion construct lacking the C-terminal RING finger required for SUMO conjugation or ubiquitination (MAPL-ΔRING-FLAG). The viruses (including an empty virus [rtTA] as negative control) were injected into the tail vein to target the expression of MAPL specifically within the liver. ABCD3 SUMOylation was restored upon expression of full length MAPL, but not in MAPL-ΔRING-FLAG (**Fig 1D**). Given the interaction between MAPL and ABCD3 observed within the BioID and IP experiments, coupled with the functional rescue of ABCD3 SUMOylation upon re-expression of MAPL, these data establish MAPL as an essential regulator of ABCD3 SUMOylation.

We next examined the consequences of the loss of MAPL on the biochemical properties of ABCD3 in liver. Consistent with our evidence that MAPL does not generally regulate protein turnover, the total mRNA and protein levels of ABCD3 were unchanged in total liver extracts as were Drp1 and Mfn2 protein levels (**Supplemental Fig 1C, D**). We also considered that peroxisomal function, biogenesis or turnover may have been globally altered in MAPL*^−/−^* liver, but observed no change in the expression levels of the peroxisomal proteins PEX14, ACOX1 and SCP2 (**Supplemental Fig 1E**).

As a half transporter, ABCD3 assembles into both homo- and heterodimers (Guimarães et al., 2004). To examine potential changes in the oligomeric assemblies of ABCD3 in the absence of MAPL, we performed sucrose gradients from solubilized mouse liver extracts. As previously described (van Roermund et al., 2014), the 70 kDa ABCD3 protein migrated at a higher molecular weight, consistent with a higher order oligomeric structure (**Fig 1E**). Consistent with the abundance of ABCD3 in the MAPL BioID, MAPL co-migrates with ABCD3 in fractions 5, 6 and 7 on the sucrose gradient. Notably, extracts isolated from MAPL*^−/−^* mice revealed a change in the migration pattern of ABCD3, with ABCD3 spreading throughout the higher molecular weight fractions, indicating that MAPL SUMOylation activity is required to maintain a stable oligomeric assembly of the ABCD3 transporter.

In sum, the data demonstrate that ABCD3 is SUMOylated by MAPL in liver tissue, a process required to regulate a peroxisomal ABCD3 complex. This provides the first evidence of a post-translational modification in the regulation of ABCD3 assembly.

### MAPL is required to gate the synthesis of bile acids in liver

To determine whether the loss of ABCD3 SUMOylation may alter circulating bile acid levels, we quantified total bile acids in plasma, and observed a 3-5-fold increase in the levels within MAPL^−/−^ mice (**Fig 2A**). Tail-vein injection of adenovirus expressing MAPL rescued the elevation in circulating bile acids in a RING-dependent manner, demonstrating the requirement for MAPL in repressing bile acid metabolism. **(Fig 2B)**. We then employed quantitative mass spectrometry using standards for over 40 selected bile acid conjugate species within circulation and in liver (see **Supplemental Fig 2A** for schematic of bile acids) (Han et al., 2015). These data revealed a significant decrease in the precursors DHCA and THCA within liver, with a concomitant increase in mature bile acid species in both plasma and liver. This indicated that the activity of bile acid synthesis was increased in these livers, leading to, for example beta-muricholic (b-MCA) acid increasing from ~10000 to 30000 fmol/g of liver tissue, and in plasma from 600 nM to 3500 nM in the MAPL*^−/−^* serum (**Fig 2C, Supplemental Fig 2B, Supplemental Table 2**).

**Figure 2:**
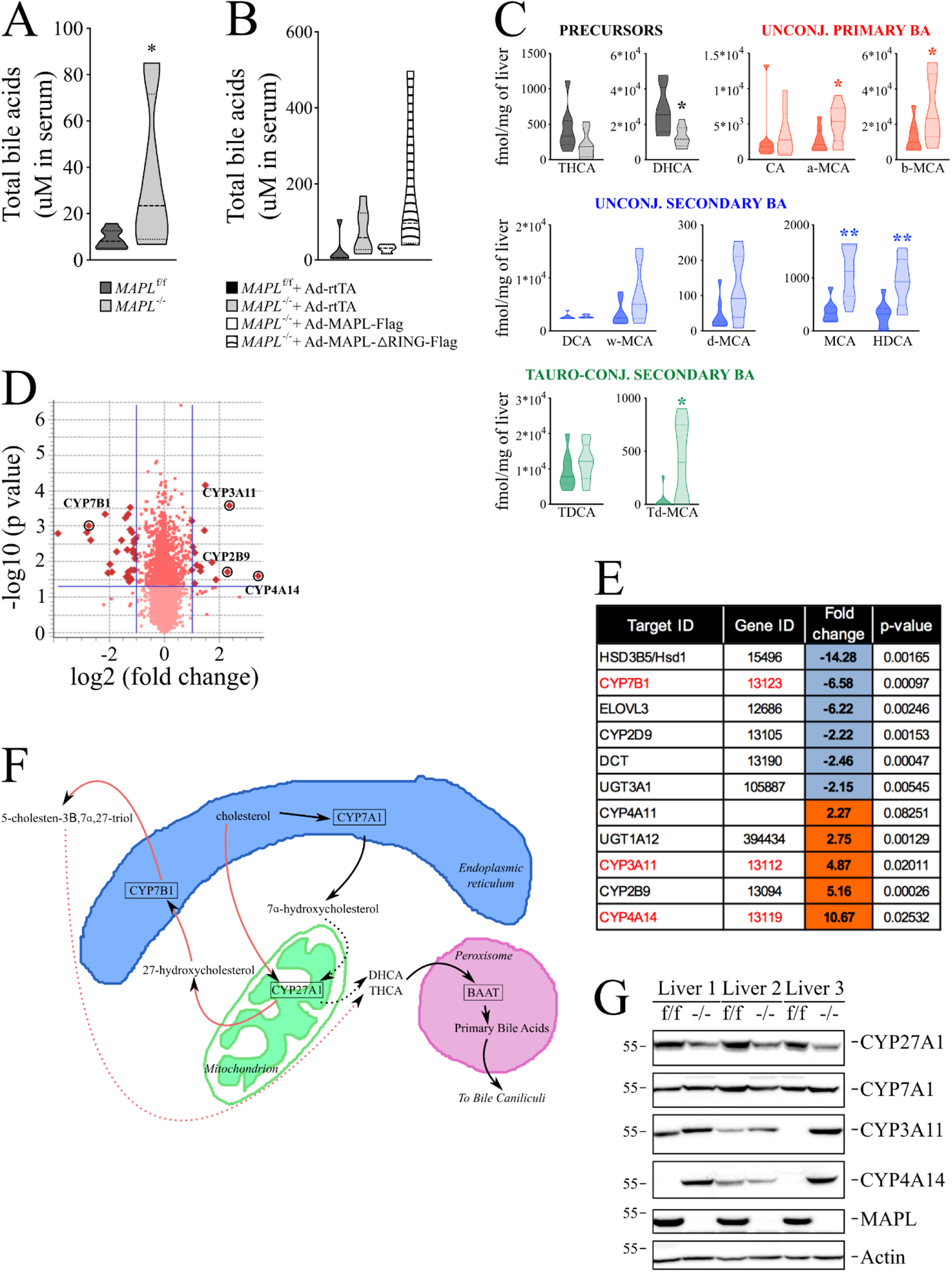
MAPL represses bile acid synthesis. **A.** Total bile acids within the serum was quantified as described in material and methods (n=8 for each strain, 6 females and 2 males, 2-6 month old). **B.** Total bile acids within the serum of tail vein injected animals was quantified as described in material and methods (n=6 with 4 females and 2 males, n=7 with 4 females and 3 males, n=4 with 2 females and 2 males and n=5 with 2 females and 3 males, for MAPL^f/f^ + rtTA, MAPL^−/−^ + rtTA, MAPL^−/−^ + MAPL-Flag and MAPL^−/−^ + MAPL-ΔRING-Flag, respectively, 2-3 month old). **C**. Bile acids precursors, as well as unconjugated and conjugated primary and secondary bile acids were quantified from liver (n=8 for each strain, 2 month old males). THCA and DHCA: tri- and dihydroxycholestanoic acid; CA: cholic acid; a-MCA, b-MCA, w-MCA and d-MCA: α-, β-, γ- and δ-muricholic acid; DCA: deoxycholic acid; MCA: murocholic acid; HDCA: hyodeoxycholic acid; TDCA: taurodeoxycholic acid; td-MCA: tauro-δ-muricholic acid. **D.** Volcano plot representation of the Illumina analysis performed on 5-month males (n=3, each strain, in triplicate). Circles highlight the significant changes in bile acid related enzymes. **E.** Table resulting from the Illumina analysis (in D) highlighting genes with variations higher than 2-fold (with a p-value <0.05, calculated with unpaired two-tailed students t-test) implicated in steroid and bile acid metabolism. **F.** A model depicting the required flux of metabolites between the ER, mitochondria and peroxisomes to facilitate bile acid synthesis in the liver. The classical pathway is represented with black arrows, while the alternative pathway is represented with red arrows. **G.** Transcriptome results were validated with western blots from whole cell liver extracts from 3 animals of each strain (one female, two males), as indicated. * *P* < 0.05 ** *P* < 0.01 in an unpaired two-tailed T test

We then performed a transcriptome analysis of liver from 8 mice of each genotype aged 5 months, revealing highly significant changes, most notably in the bile acid and steroid hormone synthesis pathways of the liver (**Fig 2D, E**, **Supplemental Table 3**). Bile acid synthesis in the liver occurs through 2 distinct pathways, the liver specific classical pathway requiring the ER localized cholesterol 7-alpha-hydroxylase, CYP7A1 (black arrows, **Fig 2F**), and the alternative or acidic pathway initiated in the mitochondria by sterol 27-hydroxylase CYP27A1 (red arrows, **Fig 2F**) that oxidizes cholesterol to 27-hydroxycholesterol (Wang et al., 2021). The 27-hydroxycholesterol derived in the mitochondria is then shuttled back to the ER where it is acted upon by 25-hydroxycholesterol 7-alpha-hydroxylase, CYP7B1 to generate 5-cholesten-3β,7α,27 triol, which is converted to the late stage C27 precursors DHCA and THCA for transport into the peroxisome. Therefore, for complete synthesis of primary bile acids, the metabolites must flux between the ER, mitochondria, peroxisomes and cytosol (**Fig 2F**). The transcriptome identified *Cyp7B1* as a major downregulated mRNA in MAPL^−/−^ liver (**Fig 2E**), however there were no available antibodies to confirm this reduction at the protein level. Therefore, we examined the protein levels of CYP27A1 and CYP7A1, representing each arm of the bile acid synthesis pathway. The data showed that the initial enzyme CYP27A1 in the alternative (or acidic) bile synthesis pathway was downregulated to 66% of control mice, consistent with a compensatory adaptation to lower total bile acid synthesis. Although liver and serum bile acids were increased significantly, the initiating enzyme of the classical pathway, CYP7A1 was unchanged (**Fig 2G, quantification Supplemental Fig 2C**). We confirmed the transcriptome data further using qRT-PCR from liver mRNA, showing a near loss of *Hsd3b5* and *Cyp7B1* mRNA, as well as a significant, 25% reduction in *Cyp27A1* mRNA (**Supplemental Fig 2D**). *Cyp7A1* and *Cyp8B1* were unchanged. In contrast, there was an upregulation of proteins involved in detoxifying bile acid intermediates and lipid soluble xenotoxins, including CYP3A11 and CYP4A14 (**Fig 2D, E, G, Supplemental Fig 2C, D**) (Wagner et al., 2005), and UDP-glucuronidation transferase 1A12 (UGT1A12, also called 1A9, **Fig 2E**), which converts lipid soluble sterols, hormones or bilirubin to water soluble, excretable metabolites (Bosma et al., 1994), all consistent with compensation to limit toxicity from elevated bile acids, and reduce synthesis. We further confirmed that MAPL depletion in liver was responsible for the near loss of *Hsd3b5* and the upregulation of *Cyp4A14*, as tail-vein injection of adenovirus expressing MAPL rescued the normal expression of these proteins in a RING-dependent manner (**Supplemental Fig 2E, F**).

We examined the established components of the regulatory feedback loop that controls *Cyp7A1* expression and found that circulating FGF15 levels were unchanged (**Supplemental Fig 2G**) (Chiang, 2009). FGF15 is secreted from the ileum in response to the bile acids that recycle across the ileum. In addition, the liver expression of the bile acid responsive Farnesoid Receptor transcription factor FXR, (gene name *Nr1h4*) was also unchanged (**Supplemental Fig 2H**) (Matsubara et al., 2013). Therefore, while we observe significant increases in bile acids in liver and plasma, the sensing system for feedback regulation of the classical pathway remained curiously unaltered.

### Increased bile acid synthesis parallels increased FGF21 secretion; that can be uncoupled from ER stress

Almost 90% of the cholesterol within the mouse liver is used to make bile, making this one of the most dominant biochemical cascades in liver (Wanders, 2013). An accumulation of intracellular bile acids have been shown to result in ER stress, leading to a transcriptional response increasing expression of the detoxifying enzymes *Cyp3A11* and *Cyp4A14* (Bochkis et al., 2008), as we observed in MAPL^−/−^ liver. An examination of ER stress markers revealed a robust increase in the phosphorylation of PERK, and CHOP expression in livers of MAPL*^−/−^* animals (**Fig 3A, quantification Supplemental Fig 3A**). Consistent with PERK activation, we observed the phosphorylation of a primary substrate, the translation initiation factor 2α, EIF2α (**Fig 3B, quantification Supplemental Fig 3A**) (Hetz, 2012). This reduces the translation of most mRNAs, allowing selective translation of the transcription factor ATF4. The upregulation of an ATF4 target gene (Salminen et al., 2017), *Fgf21* mRNA was observed in the transcriptome analysis (**Supplemental Table 3**), and by qRT-PCR we observe a robust ~30 fold increase (**Fig 3C**), and a corresponding increase at the protein level in MAPL*^−/−^* liver (**Fig 3D, quantification Supplemental Fig 3B**). *Fgf21* mRNA was increased in other tissues 2-6-fold (**Fig 3C**). Importantly, circulating levels of FGF21, quantified by ELISA showed a 12-fold increase in *MAPL^−/−^* mice (**Fig 3E**). Circulating FGF21 binds to heterotrimeric surface receptors comprised of FGFR1c, FGFR2c or FGFR3c, in complex with the β-Klotho receptors (Itoh, 2014; Owen et al., 2014). Initially thought to act primarily to signal the “browning” of white adipocytes, FGF21 binds receptors within the suprachiasmatic nucleus (SCN) of the hypothalamus and the dorsal vagal complex of the hindbrain (Bookout et al., 2013; Owen et al., 2014; Patel et al., 2015). Indeed the lean phenotype resulting from FGF21 expression was shown to be independent of the uncoupler UCP1 that is central to the browning of white adipocytes (Veniant et al., 2015). FGF21 binding within the suprachiasmatic nucleus leads to dramatic alterations in circulating glucocorticoids, altering circadian rhythm, thirst, blood pressure, and whole body metabolism (BonDurant and Potthoff, 2018; Bookout et al., 2013; Pan et al., 2018; Song et al., 2018). Consistent with these findings, we also observe a ~4-fold increase in circulating corticosterone levels, and 10-fold increase in liver of MAPL^−/−^ mice (**Supplemental Fig 3C**). In addition, evidence in rodents has shown FGF21 as a negative regulator of bile acid synthesis (Chen et al., 2018), again consistent with FGF21 upregulation within MAPL^−/−^ mice playing a potentially compensatory role to reduce bile acid synthesis.

**Figure 3:**
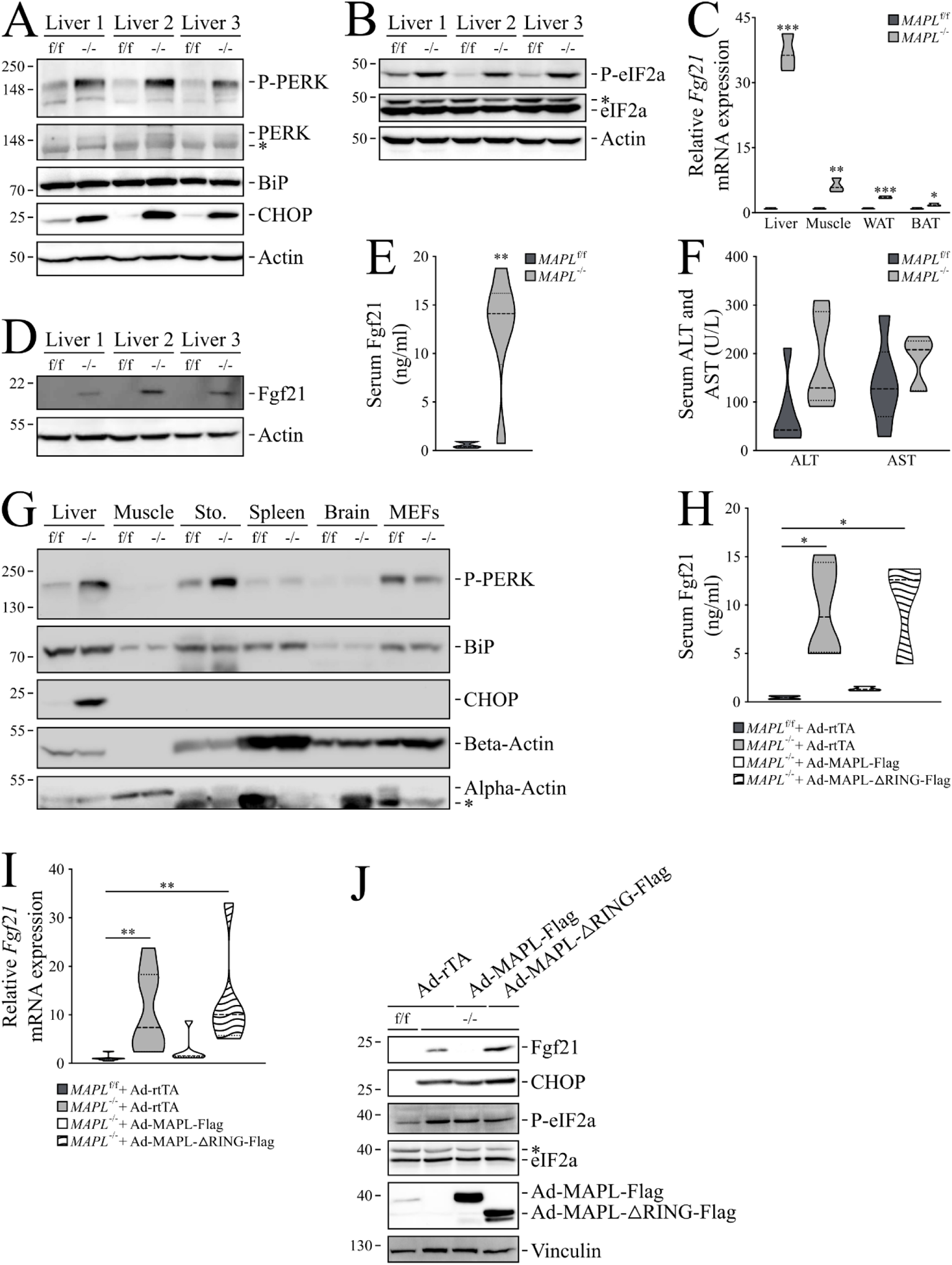
Loss of MAPL leads to hepatic ER stress, eIF2α activation and Fgf21 expression. **A.** Representative western blots from liver extracts from 3 animals of each strain: PERK auto-phosphorylation along with the total protein, as well as BIP and CHOP expression are shown (*: unspecific signal). **B.** Representative western blots of eIF2α phosphorylation in liver extracts (n=3). **C.** *Fgf21* mRNA expression measured by qRT-PCR performed on 4 different mouse tissues isolated from MAPL^f/f^ and MAPL^−/−^ animals. **D.** Representative western-blots of increased FGF21 protein levels in liver from 3 pairs of mice from each strain (left panel). **E.** Serum FGF21 was quantified by ELISA (n=8 for each strain, 2 month old males). **F.** Levels of alanine transaminase (ALT) and aspartate transaminase (AST) (n=6, 6 month old males). **G.** Representative western blots from different tissues whole cell extracts from 1 male of each strain: PERK auto-phosphorylation, as well as BIP and CHOP expression are shown (Sto: stomach, *: remaining beta actin signal). **H.** Serum FGF21 was quantified by ELISA from tail vein injected adenoviral rescued mice (n=4 with 2 females and 2 males, n=4 with 2 females and 2 males, n=3 with 3 females and n=3 with 1 female and 2 males, for MAPL^f/f^ + rtTA, MAPL^−/−^ + rtTA, MAPL^−/−^ + MAPL-Flag and MAPL^−/−^ + MAPL-ΔRING-Flag, respectively, 2-3 month old) **I.** Liver *Fgf21* gene expression by qRT-PCR on livers of rescued mice (in triplicate, n=7 with 4 females and 3 males, n=7 with 4 females and 3 males, n=5 with 3 females and 2 males and n=6 with 3 females and 3 males, for MAPL^f/f^ + rtTA, MAPL^−/−^ + rtTA, MAPL^−/−^ + MAPL-Flag and MAPL^−/−^ + MAPL-ΔRING-Flag, respectively, 2-3 month old, right panel) **J.** Representative western-blot from rescued liver extracts probed for Fgf21, CHOP, P-eIF2α and eIF2α as indicated. * *P* < 0.05 ** *P* < 0.01 *** *P* < 0.001 using T test for two group comparison and multiple comparison correction (**C,E**) or ANOVA for multiple group comparisons (**H,I**)

While there was chronic activation of ER stress within the MAPL*^−/−^* liver, the circulating levels of liver damage markers ALT and AST were only mildly increased. Therefore, ER stress did not appear pathological, and we observed no evidence of gross liver damage, steatosis or fatty liver upon histological examination (**Fig 3F, Supplemental Fig 3D, E**). As described above, any liver damage resulting from ER stress may have been ameliorated through the compensatory upregulation of detoxifying enzymes like CYP4A14 and CYP3A11 (**Fig 2G**). If the ER stress is related to increased bile acid synthesis, it should be liver specific. To test this, we examined the activation of CHOP and PERK in other tissues (**Fig 3G**). In addition to liver, we observed an increase in the phosphorylation of PERK in the stomach, but this was not accompanied by an increase in CHOP. However, there was no sign of ER stress in muscle, brain, spleen or embryonic fibroblasts, consistent with a liver specific role of MAPL in the regulation of bile acids and generation of ER stress.

As *Fgf21* is a transcriptional target of ATF4, which is selectively translated during ER (or mitochondrial) stress, it would follow that FGF21 expression should be dependent upon ER stress. However, while tail vein rescue of MAPL expression showed a complete RING-dependent restoration of circulating and liver *Fgf21* mRNA and its protein levels, the ER stress remained (**Fig 3H,I,J, Supplemental Fig 3F**). The tail-vein injection of empty adenovirus (rtTA) induced ER stress in liver, potentially masking any rescue that may have resulted from MAPL expression (**Supplemental Fig 3G**). Nevertheless, the experiment reveals an ER stress-independent regulation of FGF21 expression that instead appears to directly parallel the levels of circulating bile acids.

### MAPL^−/−^ mice are lean

A primary phenotype resulting from elevated levels of circulating FGF21 is a lean phenotype involving both autocrine and paracrine signaling pathways between the liver, brain and adipocytes (Flippo and Potthoff, 2021; Zhang et al., 2012). Consistent with these studies, mice lacking MAPL are of equal weight upon weaning, but rapidly reveal a lean phenotype, with an inability to gain weight on a high fat diet **(males in FIG 4A, B, females in Supplemental Fig 4A, B)**. Examining tissues revealed a loss in white adipocyte mass due to decreased lipid content as the primary cause of leanness, where body length and other tissue mass were unaltered **(Fig 4C, D, E, Supplemental Fig 4C, D)**. We observed browning of white fat in ~30% of mice, when examining the expression of UCP1 in white fat by both qRT-PCR and western blot analysis **(Fig 4F, Supplemental Fig 4E)**, however this did not correlate with the 100% of mice that were lean, consistent with primary targets of FGF21 in the brain. Given their metabolic phenotype, we performed insulin and glucose tolerance tests to monitor glucose handling and biogenesis. Injection of insulin led to an equivalent reduction in glycemia, however, MAPL^−/−^ mice were unable to restore their glucose levels over the 120-minute time course (males in **Fig 4G, females in Supplemental Fig 4F**). Glucose tolerance tests revealed a more rapid glucose clearance in MAPL^−/−^ vs. control mice, suggesting increased insulin sensitivity in MAPL-deficient animals (**males in Fig 4H, females in Supplemental Fig 4G**). Indeed, an analysis of circulating insulin revealed a ~3-fold reduction in insulinemia in MAPL^−/−^ compared to control mice (**Fig 4I**). In addition, glycogen storage in MAPL-deficient livers was also reduced by ~50% as compared to control livers in *ad libidum* fed animals (**Supplemental Fig 4H**). Importantly, reconstitution of MAPL expression in liver via tail vain injection increased insulinemia, thereby suggesting that the changes in insulin levels stem from the liver rather than effects in muscle, pancreas, or the periphery (**Fig 4J**). Overall, these data show that MAPL^−/−^ mice are profoundly insulin sensitive.

**Figure 4:**
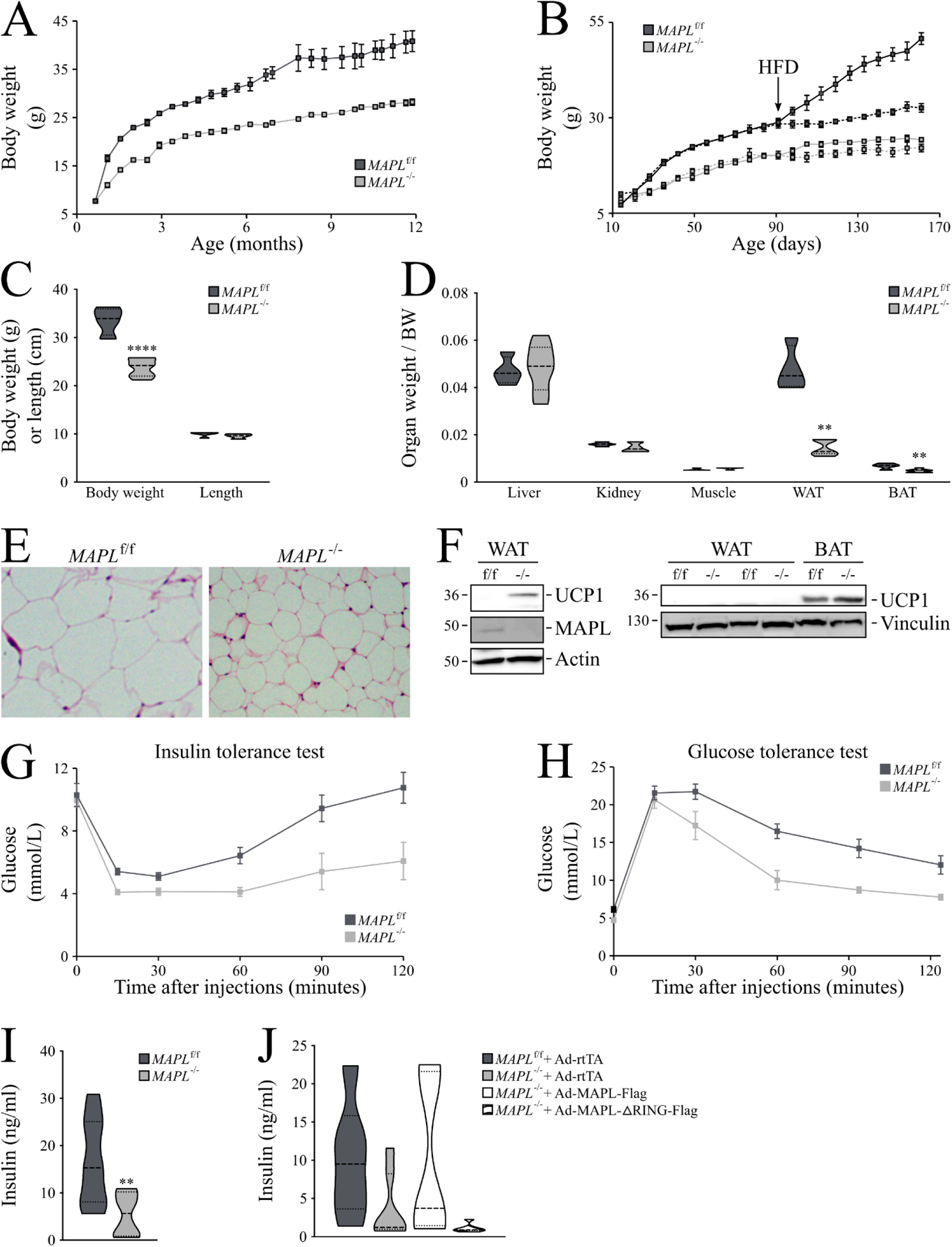
*MAPL*^−/−^ mice are lean and have increased glucose tolerance. **A.** Body weight (g) of male mice fed with normal chow; MAPL^f/f^ (n=6), and MAPL^−/−^ (n=9). **B.** Body weight (g) of MAPL^f/f^ (n=8) and MAPL^−/−^ (n=7) male mice. They were fed normal chow for 5 months (dotted lines), or for 3 months followed by 2 months of a 60% fat diet (solid lines, diet change indicated by HFD arrow). **C.** Body weight (g, left panel) or length (cm, right panel) of 7-month-old MAPL^f/f^ (n=4) or MAPL^−/−^ (n=7) male mice. **D.** Wet weight of organs including liver, kidney, gastrocnemius muscle, epididymal white fat (WAT) and interscapular brown fat (BAT) isolated from 7-month-old male MAPL^f/f^ (n=4) or MAPL^−/−^ (n=7) mice. **E.** Representative pictures of white adipocytes from MAPL^f/f^ and MAPL^−/−^ mice. Hematoxylin and eosin staining, 40X objective. **F.** Representative western-blots of Ucp1 from WAT and BAT whole cell extracts from 3 different pairs of mice (n=3). **G.** Insulin tolerance test in male MAPL^f/f^ (n=7) and MAPL^−/−^ (n=7) mice. Insulin (0.5 U/kg) was injected intraperitoneally following a 4 h fast and blood glucose was measured at indicated times. **H.** Glucose tolerance test in male MAPL^f/f^ (n=8) and MAPL^−/−^ (n=8) mice. Glucose (2 g/kg) was injected intraperitoneally following an overnight fast and blood glucose was measured at indicated times. **I.** Insulinemia (ng/mL) measured by ELISA in MAPL^f/f^ (n=8) and MAPL^−/−^ (n=8) male mice **J.** Insulinemia (ng/mL) measured by ELISA from tail vein injected adenoviral rescued mice (n=4 with 2 females and 2 males, n=4 with 2 females and 2 males, n=3 with 3 females and n=3 with 1 female and 2 males, for MAPL^f/f^ + rtTA, MAPL^−/−^ + rtTA, MAPL^−/−^ + MAPL-Flag and MAPL^−/−^ + MAPL-ΔRING-Flag, respectively, 2–3-month-old). * P < 0.05 ** P < 0.01 *** P < 0.001 **** P < 0.0001 using a repeated measures two way ANOVA and post hoc test (**G,H,I**), unpaired T test for two group comparison (**C,D**,**J**)

### MAPL^−/−^ mice develop spontaneous hepatocellular carcinoma

The loss of MAPL led to the chronic elevation of bile acids and FGF21. High levels of bile acids have been shown to promote cell proliferation and stem cell activation through TGR5 receptor binding in multiple organs (Sorrentino et al., 2020) while elevated FGF21 levels extend lifespan and improves metabolic health (Flippo and Potthoff, 2021; Zhang et al., 2012). Histology revealed atypia in the livers of MAPL^−/−^ animals, as early as 2 months, which increased in severity with age, from scattered changes to pseudo-inclusions, dysplasia, and aberrant mitotic events. (**Supplemental 5A**). This was accompanied by a ~5 and ~ 15-fold increase in hepatocyte proliferation in 2- and 7-month-old MAPL^−/−^ relative to control mice, respectively as evidenced by the Ki67 staining (**Fig 5A**). There was no obvious sign of steatosis or inflammation within MAPL^−/−^ livers, commonly linked to liver dysfunction.

**Figure 5.**
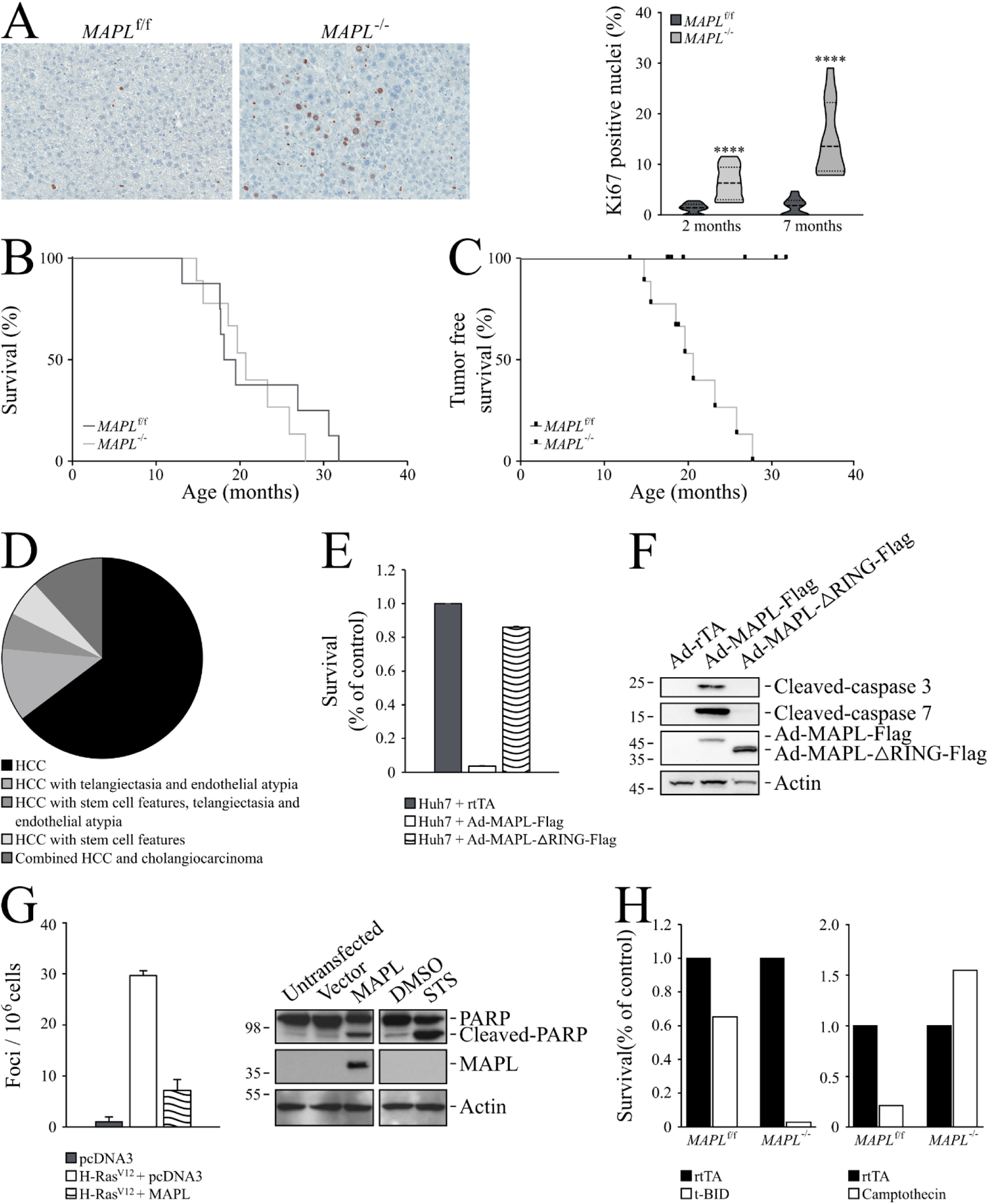
MAPL has tumor suppressive capability. **A.** Representative pictures of Ki67 liver staining (left panel) and its quantification (right panel) in MAPL^f/f^ and MAPL^−/−^ mice. **B**. Survival curve of MAPL^f/f^ (N=8) and MAPL^−/−^ (N=9) mice **C**. Cancer free survival curve of MAPL^f/f^ (N=8 and MAPL^−/−^ (N=9) male mice. **D**. Pathological analysis of liver tumors in MAPL^f/f^ and MAPL^−/−^ mice. **E**. Survival of Huh7 cells infected with adenoviruses expressing rTta, MAPL-flag, or MAPL-ΔRING-flag constructs. **F**. Immunoblots of cleaved caspase 3 and 7 from Huh7 cells infected with adenoviruses expressing rTta, MAPL-flag, or MAPL-ΔRING-flag constructs **G**. Ras foci in Rat2 fibroblasts cells transfected with empty pcDNA3 or overexpressing MAPL plasmids (left panel). Immunoblots of PARP, MAPL and beta-actin in Rat2 fibroblasts transfected with empty pcDNA3 or overexpressing MAPL plasmids (left panel). Immunoblots of PARP, MAPL and beta-actin in Rat2 fibroblasts treated with DMSO or Staurosporine (STS, 1 µM, 3 h). **H.** Survival of MAPL^f/f^ and MAPL^−/−^ primary hepatocytes infected with adenoviruses expressing rTta or truncated BID (tBID) (left panel). Survival of MAPL^f/f^ and MAPL^−/−^ primary hepatocytes treated with camptothecin (1 µM, 15 h) or vehicle. * P < 0.05 ** P < 0.01 *** P < 0.001 using T test for two group comparison and multiple comparison correction (**A**) or ANOVA for multiple group comparisons (**E,G**)

Then, we aged these animals to determine the longer-term effects of these alterations. While survival was only slightly reduced in aged MAPL^−/−^ mice relative to wild-type littermates (**males shown Fig 5B, females in Supplemental Fig 5B**), 87% and 89% (n=8 and 9) of the MAPL^−/−^ males and females, respectively, presented with liver cancer between 14-28 months (**Fig 5C, Supplemental Fig 5C**). Histological analysis from 17 tumours in different mice (both males and females) stained with H&E and reticulin confirmed all had hepatocellular carcinoma, however some of the tumours had mixed pathology (**Fig 5D**). In contrast, no littermate controls within this cohort developed any malignancies.

These findings implicate MAPL within a pathway where it exerts a tumor suppressive role in the liver. Tumor suppressors are also defined by their ability to drive or promote cell death upon overexpression. Indeed we previously described critical roles for MAPL and the SUMOylation of DRP1 in the process of apoptosis (Prudent et al., 2015), hinting that some of the cancer phenotype may arise due to an inhibition in cell death. To test whether ectopic expression of MAPL may promote cell death directly we infected a human liver cell line Huh7 with adenovirus expressing MAPL-Flag, MAPL-ΔRING-Flag SUMOylation-deficient mutant or the empty vector (Ad-rtTA). Expression of wild-type but not SUMOylation-deficient mutant MAPL led to cell death showing ~ 4% of cells remaining after 24 hours (**Fig 5E**). This was accompanied by the cleavage of caspases 3 and 7, thus indicating that forced expression of MAPL induced cell death (**Fig 5F**). Moreover, expression of SUMO competent MAPL in Rat2 fibroblasts stably expressing oncogenic H-Ras^V12^ also led to an arrest in anchorage-independent cell growth and cell death initiation, as seen with PARP cleavage (**Fig 5G**). Therefore, ectopic expression of MAPL promotes cell death and reduces neoplastic growth *in vitro*. Lastly, we examined the capacity of MAPL^−/−^ primary hepatocytes to resist cell death induced in two distinct ways. Infection of primary hepatocytes with truncated Bid (tBID) revealed that cells lacking MAPL were more sensitive to this direct induction of death. However, induction of DNA damage upon incubation with the DNA intercalating agent camptothecin showed complete resistance to cell death compared to littermate floxed hepatocytes. Therefore, while MAPL is not essential for the steps driving apoptosis downstream of activated BAX, loss of MAPL is highly protective against genotoxic stress. These data show that expression of MAPL is a driver of cell death, and loss of MAPL offers significant resistant to cell death, consistent with its liver tumor suppressor activity *in vivo*.

## Discussion

MAPL was first identified as a RING-finger containing ligase transported to peroxisomes from mitochondria in vesicular carriers, but its primary role there has been unclear. Here we present an unbiased interactome that identified previously described targets of MAPL, including Drp1 and MAVS (Braschi et al., 2009; Doiron et al., 2017; Prudent et al., 2015), which extends the list of new potential targets. This included nearly all of the machinery that regulates mitochondrial division, the ubiquitin protease USP30, and others. Previously described ubiquitin targets of MAPL were not identified in this interactome, and experiments performed in the presence of the proteasome inhibitor MG132 did not support a primary role for MAPL in the regulation of protein turnover. However, the identification of USP30 as a potential target hints that MAPL may modify USP30 directly, a deubiquitinase shown to target Parkin substrates, reversing mitophagy and Parkin-mediated protein turnover (Bingol et al., 2014; Marcassa et al., 2018). In this way, reported effects of MAPL on protein turnover and mitophagy may reflect indirect mechanisms.

While the BioID identifies multiple potential targets for MAPL, the top hit was the peroxisomal bile acid transporter, ABCD3. Our analysis of this interaction demonstrated that ABCD3 is SUMOylated in a MAPL-dependent manner *in vivo*, a modification seen to regulate the assembly into a higher molecular weight complex. This is the first documentation of SUMOylation as a regulatory post-translational modification of the bile acid transport machinery. Functionally, the loss of MAPL led to an increased production of bile acids from liver, suggesting that the SUMO conjugated form of ABCD3 may act as a gate to inhibit transport of the C27 precursors. This would be consistent with evidence for SUMO conjugation acting as a gating mechanism of potassium transporters at the plasma membrane, among others. Future work is necessary to better define how SUMOylation of ABCD3 acts to stabilize the oligomeric form and gate the channel.

The transcriptome analysis from liver also revealed a significant upregulation of *Fgf21* mRNA, which was confirmed with qRT-PCR and at the protein level in liver and plasma. Consistent with the elevation of circulating FGF21, MAPL^−/−^ mice are lean and resistant to weight gain on a high fat diet. FGF21 expression has been tightly linked to starvation and ER stress (BonDurant and Potthoff, 2018), and ER stress was observed in MAPL^−/−^ liver. However, while tail vein rescue of MAPL expression in liver demonstrated that FGF21 expression was not linked to ER stress in this system. Compensatory induction of hepatic FGF21 lead to insulin sensitivity, leanness and diet-induced obesity resistance. Finally, our data do not yet distinguish whether FGF21 expression resulted directly from the increase in bile acids through FXR (Cyphert et al., 2012) or TGR5 (Donepudi et al., 2017) signaling, or whether it may relate to an unknown functional target of MAPL in liver.

An interesting aspect of this study is that loss of MAPL led to an alteration in the “alternative” pathway of bile acid synthesis without any change in the canonical, classical pathway. Although the transcriptional regulation of the classical enzyme CYP7A1 is very well studied (Chiang, 2009), the regulation of the more broadly expressed CYP27A1 and CYP7B1 is less clear (Stiles et al., 2009). The latter enzymes are expressed in multiple tissues, playing roles in sterol conversion and cholesterol homeostasis in different cell types. The regulation of CYP27A1 expression is tissue specific and broad, with links to bile acid feedback, PPAR agonists, insulin signaling, growth hormones and glucocorticoids (Lorbek et al., 2012). A recent study showed a specific upregulation of CYP7B1 during cold exposure, leading to increased bile acid secretion, altered microbiome and heat production (Worthmann et al., 2017). The mechanisms regulating CYP7B1 expression in that study was not elucidated and we now reveal a role for MAPL in modulating this specific arm of bile acid metabolism. In the case of MAPL^−/−^ mice, the reduction in CYP7B1 and CYP27A1 appear to be compensatory, accompanying increases in sterol clearance and detoxification pathways, likely minimizing liver damage. Circulating bile acids have been shown to bind to the G-coupled protein receptor TGR5 within multiple organs including adipocytes, brain, and gut (de Boer et al., 2018). Therefore, the global phenotype of MAPL deficient mice will almost certainly be impacted by the elevated circulating bile in multiple ways, along with potential cell autonomous functions of MAPL in different tissues. Our study has focused first on the primary phenotype in liver where our study identified a critical function for MAPL in bile acid metabolism.

Ultimately, mice lacking MAPL also showed increased hepatocyte proliferation and later development of hepatocellular carcinoma. While this is consistent with emerging roles for bile acids driving hepatocyte proliferation (Anakk et al., 2013), the circuitry of these events will also be a focus of our future work.

MAPL/MUL1 has been previously linked to numerous cellular processes including mitophagy (Ambivero et al., 2014; Li et al., 2015; Rojansky et al., 2016; Yun et al., 2014), inflammation (Barry et al., 2018; Ni et al., 2017), antiviral (Doiron et al., 2017; Jenkins et al., 2013), apoptosis (Prudent et al., 2015) and proliferation (Jung et al., 2011; Zhang et al., 2008), where its activity allows the dynamic and rapid regulation of diverse signals. Our previous studies demonstrated DRP1 as another substrate of MAPL, playing a key role in stabilizing the oligomeric DRP1 during cell death. The fission machinery was identified within the BioID here as well and we demonstrate that ectopic expression of MAPL activated cell death pathways. Our data also reveal that loss of MAPL restrict cell death pathways induced by genotoxic stress, that may further contribute to tumor formation. Our future work will continue to investigate how the tumor suppressive activity of MAPL functions in the regulation of global metabolism in liver, and the potential relevance of MAPL function within human cancers. The *MUL1* gene lies on chromosome 1p36, which is a very commonly deleted region in human cancer. While there are many genes within this region of the chromosomes, the NCI cancer genome atlas reports the most common cancer with loss of *MUL1* is cholangiocarcinoma. For now, the MAPL^−/−^ mice provide a new model to better understand the complex signaling pathways within in different tissues, and under a variety of stimuli.

## Methods

### Ethics Approval

Animal experimentation was conducted in accordance with the guidelines of the Canadian Council for Animal Care, with protocols approved by the Animal Care Committees of the University of Ottawa and of McGill University.

### Generation of floxed MAPL KO mice

The targeting vector and the *MAPL^WT/flox^* mice were generated by Ozgene (Australia). The construct contained two loxP sequences inserted in intron 1 and intron 2 of the *MAPL* gene, and two frt sites flanking the neomycin resistance selection cassette. The construct was electroporated into C57BL/6 ES cell line, Bruce4 (Köntgen et al., 1993). Homologous recombinant ES cell clones were identified by Southern hybridization and injected into BALB/cJ blastocysts. Male chimeric mice were obtained and crossed to C57BL/6J females to establish heterozygous germline offspring on pure C57BL/6 background. To remove the Neo-cassette (neo), *the MAPL^WT/flox^* mice were bred with homozygous FlpE-“deletor” C57BL/6 mice (Ozgene).

To generate *MAPL^−/−^* mice, *MAPL^f/f^* mice were first bred with *CMV-Cre* carrying out mice (The Jackson Laboratory). The resulting *MAPL^+/-:Cre^* were then bred with *MAPL^f/f^* animals. One quarter of the offspring were *MAPL^f/-^*. These heterozygous mice, devoid of the *CMV-Cre* gene, were used as breeders: their offsprings were composed of 25% *MAPL^−/−^* animals, 25% of *MAPL^f/f^* animals used as littermate wild type controls and 50% of *MAPL^f/-^* animals used as littermate heterozygous controls.

Genotyping was performed by PCR of tail DNA (extracted using the DNA Blood & Tissue kit, QIAgen, according to the manufacturer’s instruction) using two different primer pairs (Primer1: Fwd :5’-GGGAAGTGTGTGCCTTATG Rev: 5’-AATCCCAAGTCCACAGTGC and Primer2: Fwd: 5’-CCTCAGAGTTCATTTATCC Rev: 5’-CCAACACCATCAAAAGGC).

Mice were fed *ad liditum* either normal chow or a high fat diet (60% fat, 20% proteins, 20% carbohydrate, Research Diets, for 10 weeks, starting at 12 weeks old). The food intake and body weight of each mouse were recorded weekly.

### Metabolic tests

Glucose tolerance tests (GTT) were performed after an overnight (16h fast). Blood glucose and plasma insulin levels were measured after intra-peritoneal injection of glucose (2 g/kg of body weight). Insulin tolerance tests (ITT) were performed after intra-peritoneal injection of human insulin (0.5 U/kg) in 2-h-fasted mice.

### Primary hepatocytes isolation, culture and glucose production

Primary hepatocytes were isolated from 12- to 16-week-old mice by 2-step liberase perfusion (Liberase TL; MilliporeSigma #05401020001) and 50% Percoll gradient purification (MilliporeSigma #P1644). Cells were plated on collagen coated plates and cultured in Dulbecco’s modified Eagle’s medium supplemented with 0.2% bovine serum albumin (fatty acid free; Fisher Scientific), 25 mM glucose, 2 mM sodium pyruvate, 0.1mM dexamethasone, 1% penicillin/streptomycin, and 1 nM insulin for up to 48 hours. To measure glucose production, primary hepatocytes were switched to basic medium (DMEM with 0.2% BSA and 1 mM glutamine, with no glucose, red phenol or sodium pyruvate) for 2 h to induce glycogenolysis and deplete glycogen. Basic media containing 2 mM lactate, 1 mM pyruvate, 1 mM glycerol with or without 10 nM glucagon (to promote gluconeogenesis) was exchanged and harvested every hour for 3 h. Glucose released was measured by enzymatic reaction (Hexokinase assay #GAHK20 MilliporeSigma) and normalized to protein content per well.

### Adenovirus tail vein injection

2*10^9^ PFU/mouse (MAPL-Flag) or 0.67*10^9^ PFU/mouse (MAPL-ΔRING-Flag and rtTA) of adenoviruses were injected through the tail vein in a 100 µl final volume of sterile saline solution to 2-3 month old animals. At day 7 post injection, mice were starved overnight and fed back for 3 hours (from 8 to 11AM). Blood was then collected by cardiac puncture and livers were collected for further investigations.

### Electrophoresis and immunoblot analysis

Tissues were homogenized in ice-cold lysis buffer (40 mM NaCl, 2 mM EDTA, 1 mM orthovanadate, 50 mM NaF, 10 mM pyrophosphate, 10 mM glycerolphosphate, 20 mM NEM, 1% Triton X-100, 50 mM Hepes, pH 7.4) supplemented with Complete protease inhibitor cocktail (Roche Molecular Biochemicals) in a borosilicate glass Dounce tissue grinder with tight pestle. After 20 min at 4°C, homogenates were centrifuges at 20,000 g for 20 min at 4°C, and the supernatants were collected. Protein extracts (20 µg) were separated on a Tris-Glycine 4-20% gradient precast polyacrylamide gel (Invitrogen), and transferred to 0.22 µm pore nitrocellulose membrane (Bio-Rad). Bands were visualized with Western-Lightning PLUS-ECL (Perkin-Elmer) with an INTAS ChemoCam (INTAS Science Imaging GmbH) and quantified with ImageJ software

MAPL was detected by rabbit polyclonal antibodies (HPA017681, 1:1,000, Sigma), PERK by rabbit polyclonal antibodies (100-401-962, 1:1,000, Rockland antibodies & assays), phospho-PERK by rabbit monoclonal antibodies (3179, 1:1,000, Cell Signaling), eIF2α by mouse monoclonal antibodies (2103, 1:500, Cell Signaling), phospho-eIF2α by rabbit polyclonal antibodies (SAB4504388, 1:500, Sigma), Fgf21 by goat polyclonal antibodies (AF3057, 1:500, R&D systems), CHOP by rabbit polyclonal antibodies (5554, 1:500, Cell Signaling), BiP by rabbit polyclonal antibodies (ADI-SPA-826-D, 1:1000, Enzo), CYP3A11 by rabbit polyclonal antibodies (13384, 1:500, Cell Signaling), CYP4A14 by goat polyclonal antibodies (sc-46087, 1:500, Santa Cruz), CYP7A1 by rabbit polyclonal antibodies (ab65596, 1:500, Abcam), CYP27A1 by rabbit polyclonal antibodies (NBP2-16061, 1:500, Novus Biologicals), SUMO1 by mouse monoclonal antibodies (332400, 1:1000, Invitrogen), Hsp60 by mouse monoclonal antibodies (sc-136291, 1:1000, Santa Cruz), Hsp70 by rabbit polyclonal antibodies (ab137680, 1:1000, Abcam), ABCD3 by mouse monoclonal antibodies (sab4200181, 1:1000, Sigma), DRP1 by mouse monoclonal antibodies (611113, 1:1000, BD Transduction Labs), Mfn2 by rabbit polyclonal antibodies (M6319, 1:1000, Sigma), UCP1 by a polyclonal antibody (U6382, 1:500, Sigma), ACOX1 by a polyclonal antibody (10957-1-AP, 1:1000, Proteintech), SCP2 by a polyclonal antibody (14377-1-AP, 1:1000, Proteintech), PEX14 by a polyclonal antibody (ABC142, 1:1000, Millipore), vinculin by a monoclonal antibody (V4505, 1:1000, Sigma), β-actin by rabbit polyclonal antibodies (SAB4502543, 1:1,000, Sigma) and β-actin by mouse monoclonal antibodies (A2228, 1:1,000, Sigma).

### Cellular fractionation

Liver was collected into ice-cold PBS and rinsed free of blood. It was minced into small pieces and homogenized using a Dounce homogenizer (3-4 times, 1,600 rpm) into IB isolation buffer (mannitol 200 mM, sucrose 68 mM, Hepes 20 mM pH 7.4, KCl 80 mM, EGTA 0.5 mM, Mg(Acetate)2 2 mM, 2-chloroacetamide 20 mM and protease inhibitors 1X). Homogenate was centrifuged at 800g for 10 min to separate nuclear pellet from post-nuclear supernatant. The nuclear pellet was resuspended into 2 ml of IB buffer and centrifuged once again at 800 g for 10 min. Pellet was resuspended into 200 µl IB and kept as nuclear fraction. The post nuclear supernatant was centrifuged at 1,000 g for 10 min. The supernatant was kept and centrifuged at 10,000 g for 20 min to separate mitochondrial heavy membrane pellet and post-mitochondrial supernatant. Mitochondrial pellet was washed in 1 ml IB and centrifuged at 10,000 g for 10 min. The final mitochondrial pellet was resuspended into 50 µl IB and kept as mitochondrial fraction. The post-mitochondrial supernatant was centrifuged at 200,000 g in TLA-110 rotor (Beckman-Coulter) for 40 min. Supernatant was kept as cytosolic fraction.

### Immunoprecipitation

For the MAPL immunoprecipitation, livers from 4 month old males were washed in ice-cold PBS and homogenized in 5ml of lysis buffer (50 mM Tris, 150 mM NaCl, 0.5 mM EDTA, 2 mM MgCl^2^, 1% triton X-100, 20 mM NEM, pH 7.5) supplemented with Complete protease inhibitor cocktail (Roche Molecular Biochemicals) in a borosilicate glass Dounce tissue grinder with tight pestle. After 20 min at 4°C rocking, homogenates were centrifuged at 20,000 g for 20 min at 4°C and supernatants were collected. 1mg of proteins (diluted at 2 mg/ml in lysis buffer) was pre-cleared overnight at 4°C, rocking with 100 µl of Dynabeads protein A beads (Life Technologies). 100 µl of Dynabeads protein A beads (resuspended in 200 µl of 0.1 M NaP, 0.08% Tween 20) per condition were incubated overnight with 5 µg of antibodies. The next day, antibodies were covalently bound to the beads using DMP crosslinker (Pierce) 20 mM, for 30 min at RT, rocking in the dark) and crosslink reaction was stopped by 50 mM Tris pH 7.5 for 15 min at RT, rocking. Beads were washed 2 times with 100 µl of 0.1 M glycine pH 2.5. Precleared homogenates (1:20 was saved as starting material, SM) were applied on the antibodies-bound beads overnight at 4°C, rocking. The next day, homogenates were removed from the beads and beads were washed 2 times with lysis buffer, 2 times with high salt buffer (50 mM Tris, 450 mM NaCl, 0.5 mM EDTA, 2 mM MgCl^2^, 0.05% triton X-100, 20 mM NEM, pH 7.5) and 2 times with low salt buffer (50 mM Tris, 150 mM NaCl, 0.5 mM EDTA, 2 mM MgCl^2^, 0.05% triton X-100, 20 mM NEM, pH 7.5). Proteins were eluted with 50 µl of 0.1 M glycine, pH 2.5, 0.5% Triton X-100 (2 times). 20 µl of Tris 1 M pH 7.5 were added to the elution.

### SIM-beads extracts from livers

Mouse livers were washed in ice-cold PBS and resuspended in cell fractionation buffer. Cells were then broken with a cell cracker (EMBL-Heidelberg) using ball size 8.002. The samples were centrifuged at 800 g at 4°C for 10 min. Supernatants were re-cleared at 800 g for 5 min at 4°C. Post-nuclear supernatants were then centrifuged at 9,000 g for 20 min at 4°C, in order to pellet heavy membrane fraction (supernatant is the cytosolic fraction). The pellets were washed and re-centrifuged for a further 10 min at 9,000 g. To heavy membrane and cytosolic fractions, Triton X-100 was added up to 1% concentration, followed by incubation for 20 min at 4°C, rocking. The fraction lysates were then centrifuged for 45 min at 200,000 g at 4°C. Supernatants were then collected and protein concentration determined (80 µg total lysate fraction of each type were separated to run in gel as starting material). Then 40 µl of SIM beads (AM-200, Boston Biochemicals) were added to each type of total lysate fractions (containing 0.5-1 mg of proteins) and incubated for at least 1.5 hours at 4°C, rocking, followed by centrifugation at 14,000 g for 2 min at 4°C. Beads were then washed 5 times with cell fractionation buffer containing 1% Triton X-100. After the last wash was discarded, 1X Laemmli sample buffer was added to the solid beads and run in acrylamide gel, together with the starting material.

### BirA and Flag-MAPL-BirA stable cells

HEK293T-REX cells stably expressing either BirA or Flag-MAPL-BirA were maintained in DMEM (Wisent) containing 10 % FBS (Wisent), 2 mM L-glutamine, non-essential amino acids and 1 mM sodium pyruvate (Life Technologies). The expression of BirA and Flag-MAPL BirA was induced with 1 ug/ml tetracycline (Sigma) for 24 hours in culture at 37°C, accompanied by 50 µM biotin (Sigma) final concentration. After the incubation, the cells on plates were washed 3 times in PBS, then they were scrapped on ice and lysed for 20 min in buffer: 10 mM Hepes pH 7.4, 200 mM NaCL, 0.5 mM EDTA, 2 mM MgCL2, 1 % Triton-X100. Lysates were centrifuged (20,000 g at 4°C for 15 min), normalized for protein concentration and incubated with streptavidin-agarose beads (Life Technologies) for 1.5 hour rocking at 4°C. Beads were centrifuged, washed 3 times with lysis buffer, mixed with 1 X Laemmli buffer, and loaded onto a gel, along with starting material. Then electrophoresis was performed, and proteins were transferred to nitrocellulose membranes. Expression of Flag-BirA-MAPL, as well other candidate proteins for interacting with MAPL, was determined by western blot with a series of antibodies.

### Sucrose gradient

Livers from 2-month-old males were washed in ice-cold PBS and homogenized in 3 ml of homogenization buffer (50 mM Tris, 150 mM NaCl, 0.5 mM EDTA, 2 mM MgCl^2^, 20 mM NEM) supplemented with Complete protease inhibitor cocktail (Roche Molecular Biochemicals). A fraction of the homogenates was sonicated in order to quantify protein content. Homogenates were diluted in homogenization buffer to 4.7 mg/ml. 500 µl of homogenate at 4.7 mg/ml were added to 500 µl of homogenization buffer containing 1% DDM. After 20 min at 4°C rocking, homogenates were centrifuged 20 min at 4°C at 14,000 rpm. Supernatants were collected, and protein content determined. 250 µg of proteins were loaded on the top of the 10-50% sucrose linear gradient (1800 µl, 200 µl of each) and centrifuged for 4 hours at 4°C, at 180,000 g. 12 fractions were collected and 75µ l of the different fractions were loaded on a 10% acrylamide gel.

### Blood analysis

Each tube was added with 250 µL acetonitrile and the samples were homogenized again, followed by centrifugation at 15000 rpm and 10°C for 5 min. 200 µL of the supernatants were mixed with 50 µL of the same IS solution, followed by PD-SPE using the same procedure as done for mouse serum. The residues were reconstituted in 200 µL of 50% methanol.

20 µL of each of the above samples was injected onto a 15-cm long C18 UPLC column for quantitation of bile acids by UPLC-(-)ESI/MRM/MS with negative-ion mode detection and with water-acetonitrile-formic acid as the mobile phase for binary gradient elution, using the same method as described in the publication. UPLC-MRM/MS runs were performed on a Dionex Ultimate 3000 UPLC system coupled to a 4000 QTRAP triple-quad mass spectrometer.

Concentrations of the detected bile acids were calculated with internal standard calibration from calibration curves prepared for individual compounds. For concentration calculation, the 14 D-labeled bile acids were used as IS for their corresponding non-D-labeled forms. For the bile acids, THCA and DHCA, for which there were no D-labeled analogues as IS, chenodeoxycholic-D4 acid was used as the common IS for quantitation of the unconjugated bile acids, THCA and DHCA; tauro-CDCA-D4 was used as the common IS for quantitation of the taurine-conjugated species; glyco-deoxycholic-D4 acid was used as the common IS for quantitation of the glycine-conjugated species.

Concentrations of the following bile acids were also estimated in this analysis: glyco-ω-MCA, glyco-α-MCA, glyco-β-MCA, glyco-λ-MCA (also as glyco-γ-MCA or glycohyocholic acid) or glyco-allocholic acid. Since there were no standard substances for these compounds, their concentrations were calculated from the calibration curve of glycocholic acid.

Another 20 µL of each of the same samples was injected again onto the same C18 UPLC column for UPLC-MRM/MS quantitation of corticosterone, but with positive-ion (+) mode detection.

### Illumina

Total RNAs from the liver of 3 mice from each strain (5 months old) were isolated using the TRIzol kit, following manufacturers protocols (Invitrogen), as described bellow. RNAs were quantified using a NanoDrop Spectrophotometer ND-1000 (NanoDrop Technologies, Inc.) and its integrity was assessed using a 2100 Bioanalyzer (Agilent Technologies). Double stranded cDNA was synthesized from 250ng of total RNA, and *in vitro* transcription was performed to produce biotin-labeled cRNA using Illumina® TotalPrep RNA Amplification Kit, according to manufacturer’s instructions (Life Technologies). The labeled cRNA was then normalized at 1500ng and hybridized on Mouse WG-6, v.2 according to Illumina’s Whole-Genome Gene Expression Direct Hybridization Assay Guide. The BeadChips were incubated in an Illumina Hybridization oven at 58°C for 14 to 20 hours at a rocking speed of 5. Beadchips were washed also according to Illumina’s Whole-Genome Gene Expression Direct Hybridization Assay Guide and scanned on an Illumina iScan Reader. RNA from each mouse was sequenced in triplicate. Results were analyzed using the FlexArray software (provided by Genome Quebec). Volcano plot was calculated using a 2 fold increase and decrease lower limit, with p values lower or equal to 0.05.

### BioID

BioID (Roux et al., 2012) was carried out essentially as described previously (Comartin et al., 2013). In brief, the full-length human MAPL (BC014010) coding sequence was amplified by PCR and cloned into a pcDNA5 FRT/TO BirA*FLAG expression vector (MAPL-AscI_Fwd: tataGGCGCGCCaATGGAGAGCGGAGGGCGGCCCTCG; MAPL-NotI_Rev: ttaaGCGGCCGCGCTGTTGTACAGGGGTATCACCCG). Using the Flp-In system (Invitrogen), 293T-REx Flp-In cells stably expressing MAPL-BirA*Flag were generated. After selection (DMEM + 10% FBS + 200 µg/ml hygromycin B), 10 x 150 cm^2^ plates of subconfluent (60%) cells were incubated for 24 hours in complete media supplemented with 1 µg/ml tetracycline and 50 µM biotin. Five plates were treated with 5 µM MG132. Cells were collected and pelleted (2000 rpm, 3 min), the pellet was washed twice with PBS, and dried pellets were snap frozen. Pellets were lysed in 10 ml of modified RIPA lysis buffer (50 mM Tris-HCl, 150 mM NaCl, 1 mM EDTA, 1 mM EGTA, 1% Triton X-100, 0.1% SDS, 1:500 protease inhibitor cocktail, 250U Turbonuclease, pH 7.5) at 4°C for 1 hour, then sonicated to completely disrupt visible aggregates. The lysates were centrifuged at 35,000 g for 30 min. Clarified supernatants were incubated with 30 µl packed, pre-equilibrated Streptavidin-sepharose beads at 4°C for 3 hours. Beads were collected by centrifugation, washed 6 times with 50 mM ammonium bicarbonate pH 8.2, and treated with TPCK-trypsin (16 hours at 37C). The supernatant containing the tryptic peptides was collected and lyophilized. Peptides were resuspended in 0.1% formic acid and 1/6^th^ of the sample was analyzed per MS run.

Liquid chromatography (LC) analytical columns (75 µm inner diameter) and pre-columns (100 µm ID) were made in-house from fused silica capillary tubing from InnovaQuartz and packed with 100Å C^18^-coated silica particles. LC-MS/MS was conducted using a 120 min reversed-phase buffer gradient running at 250 nl/min (column heated at 35C) on a Proxeon EASY-nLC pump in-line with a hybrid LTQ-Orbitrap Velos mass spectrometer. A parent ion scan was performed in the Orbitrap, using a resolving power of 60000. Simultaneously, up to the 20 most intense peaks were selected for MS/MS (minimum ion count of 1000 for activation) using standard CID fragmentation. Fragment ions were detected in the LTQ. Dynamic exclusion was activated such that MS/MS of the same *m/z* (within a 10 ppm window, exclusion list size 500) detected 3 times within 45 sec were excluded from analysis for 30 sec. For protein identification, .RAW files were converted to the mzXML format using Proteowizard, then searched using X!Tandem against human RefSeq Version 45 (containing 36113 entries). Search parameters specified a parent MS tolerance of 15 ppm and an MS/MS fragment ion tolerance of 0.4 Da, with up to 2 missed cleavages allowed for trypsin. Oxidation of the methionine was allowed as a variable modification. Data were analyzed using the trans-proteomic pipeline via the ProHits 2.0.0 software suite. Proteins identified with a ProteinProphet cut-off of 0.85 (corresponding to ≤1% FDR) were analyzed with SAINT Express v.3.3. Sixteen control runs were used for comparative purposes, comprising 8 runs of BioID conducted on untransfected 293T-REx cells. In each case, 4 runs were conducted on untreated cells, and 4 runs were conducted in cells treated with MG132, as above. The 16 controls were collapsed to the highest 4 spectral counts for each hit. All raw mass spectrometry data have been uploaded to the MassIVE archive (ucsd.edu), ID: MSVxxxx, password: MAPL.

### Histology

Formaldehyde-fixed, paraffin-embedded tissues were cut into 4µm sections and stained with hematoxylin and eosin (H&E).

### RNA isolation and qRT-PCR

Total RNAs from various tissues were prepared using TRIzol (Invitrogen). They were treated with DNAse (New England Biolabs), then reverse transcribed with random primers using the High Capacity sDNA Reverse Transcription Kit (Life Technologies) as described by the manufacturer. Before use, RT samples were diluted 1:5. Gene expression was determined using assays designed with the Universal Probe Library (UPL) from Roche (www.universalprobelibrary.com). For each qPCR assay, a standard curve was performed to ensure the efficacity of the assay is between 90% and 110%. qPCR reactions were performed using 5-25 ng of cDNA samples, the TaqMan Advanced Fast Universal PCR Master Mix (Life Technologies), 2 µM of each primer and 1µM of the corresponding UPL probe. The Viia7 qPCR instrument (Life Technologies) was used to detect the amplification level and was programmed with an initial step of 3 min at 95°C, followed by 40 cycles of: 5 sec at 95°C and 30 sec at 60°C. All reactions were run in triplicate and the average values of Cts were used for quantification. The relative quantification of target genes was determined using the ΔΔCT method. Briefly, the Ct (threshold cycle) values of target genes were normalized to an endogenous control gene (ΔCT = Ct ^target^ – Ct ^CTRL^) and compared with a calibrator: ΔΔCT=ΔCt^Sample^-ΔCt^Calibrator^. Relative expression (RQ) was calculated using the Sequence Detection System (SDS) 2.2.2 software (Applied Biosystems) and the formula is RQ = 2^−ΔΔCT^.

### qRT-PCR primers used (5’ to 3’)

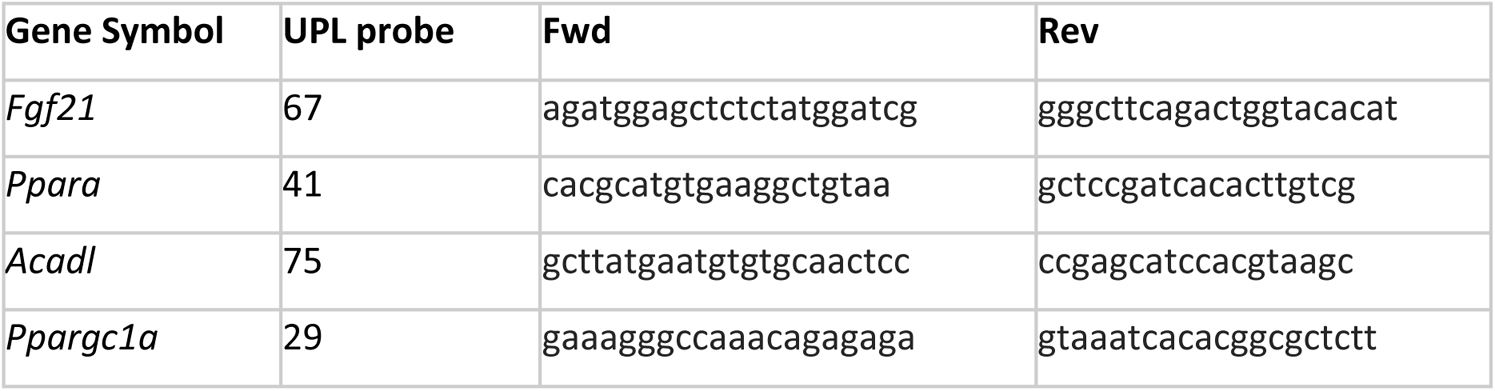
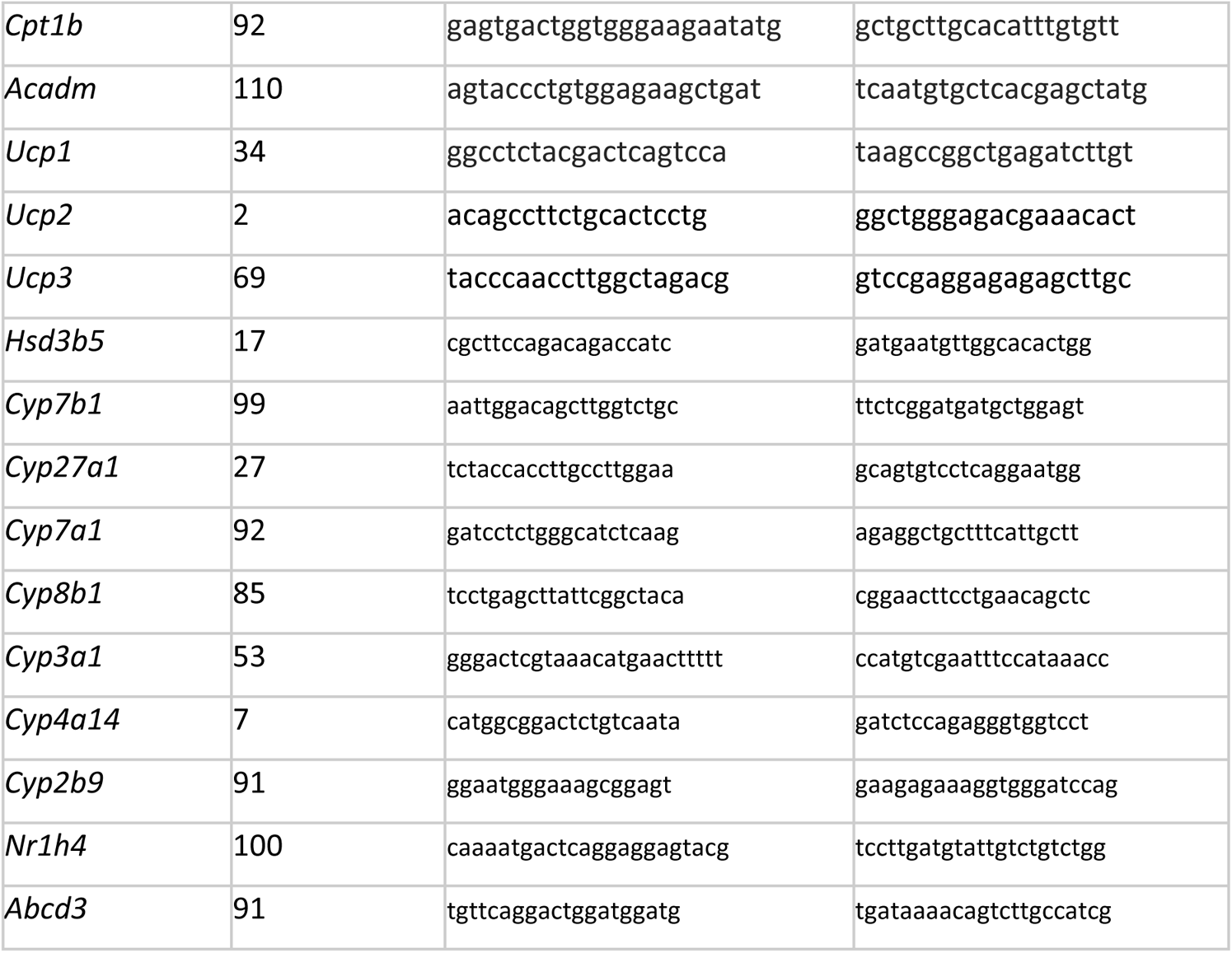

### Statistical analysis

Normal distribution and homoscedasticity of data were tested by Shapiro-Wilks and Bartlett tests respectively. Parametric tests were used if distributions normal and variances equal. Student t-test was used to compare 2 groups on 1 variable. One-way analysis of variance (ANOVA) for univariate multiple comparisons or Two-way ANOVA (for bivariate comparisons) were followed by Tukey’s honest significant difference post-hoc test. Statistical analyses were performed using GraphPad Prism software (San Diego, CA). Threshold for statistical significance was *P* < 0.05.

All values are expressed as mean ± SEM

## Acknowledgements

This study was funded by Canadian Institute of Health Research and the Canadian Cancer Society Research Institute (to HMM; CIHR#68833, CCSRI#702139). Work in the BR lab was funded by CIHR #130340. HMM holds the Canada Research Chair in Mitochondrial Cell Biology. BR holds the Canada Research Chair in Proteomics and Molecular Medicine. ABP was supported by a postdoctoral fellowship from Fonds de Recherche Québec - Santé. The authors would like to thank the IRIC for qRT-PCR services and Ozgene (Australia) for services in the design and generation of the floxed MAPL strain. We also thank Dr. Atilla Omeroglu (McGill University) for help with histological analysis, Nancy Braverman (McGill University) for advice on bile acid metabolism, and Tharan Srikumar for mass spectrometry technical assistance and the MNI animal care services.

## Figures

**Figure S1:**
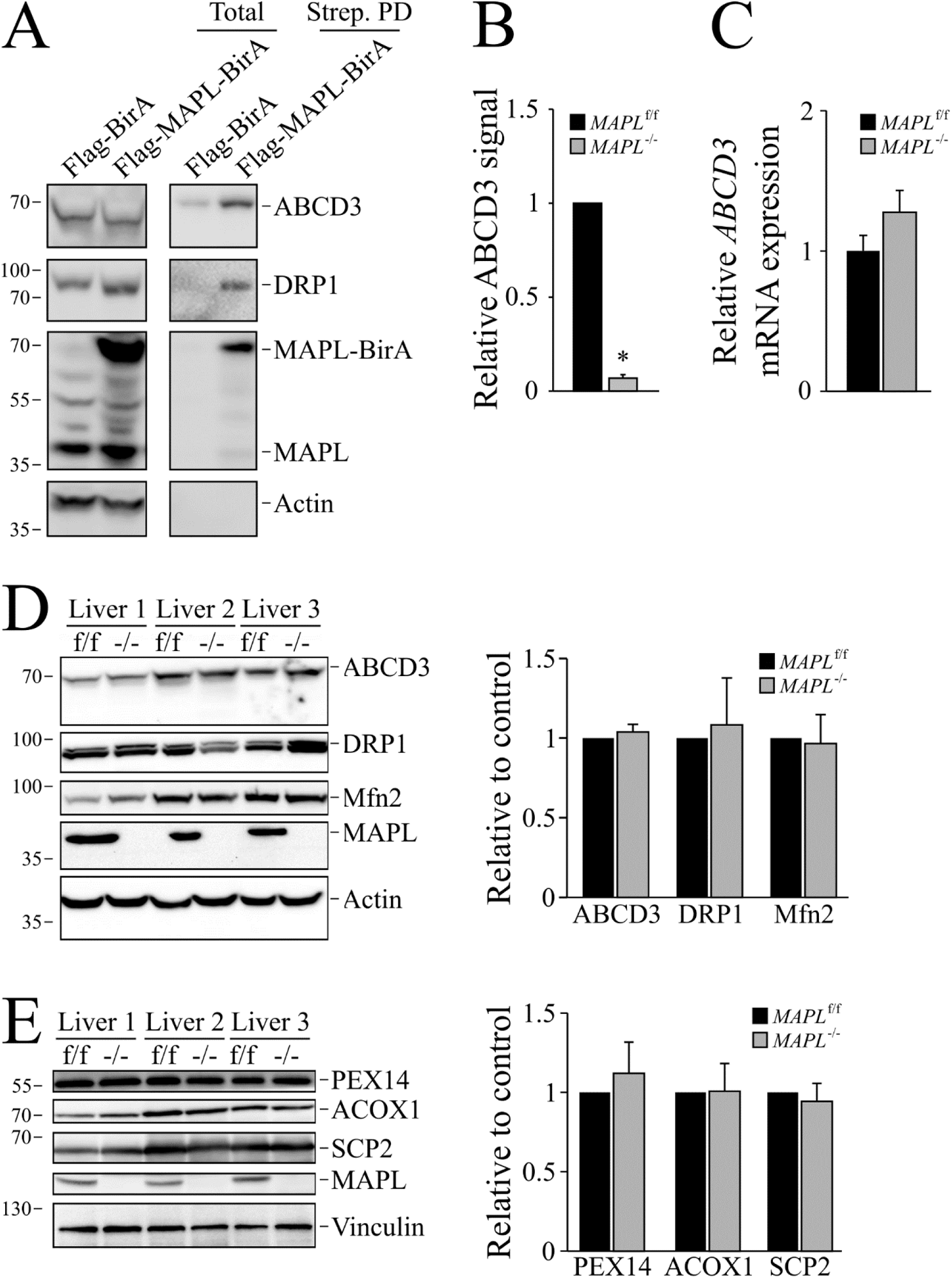
MAPL SUMOylates ABCD3 and its absence alters ABCD3 complex assembly. **A.** Representative western-blot of total lysate and streptavidin-beads pull-down fractions of HEK-293T-REX cells expressing Flag-MAPL-BirA or Flag-BirA (as control) incubated with biotin and probed for ABCD3, DRP1, MAPL and beta-actin. **B.** Quantification from 3 independent experiments of ABCD3 signals of the heavy membrane SIM-beads elution fraction (Figure 1C). **C.** qRT-PCR investigating *ABCD3* mRNA levels in livers of 4 different mice (2-month-old males). **D.** Representative western blots from whole cell liver extracts probed for ABCD3, DRP1 and Mfn2 (Left panel). ABCD3, Drp1 and Mfn2 signals were quantified (n=3 for each strain, right panel). **E.** Representative western blots from whole cell liver extracts probed for PEX14, ACOX1 and SCP2 (Left panel). PEX14, ACOX1 and SCP2 signals were quantified (n=3 for each strain, right panel). * *P* < 0.05 ** *P* < 0.01 *** *P* < 0.001 using T test for two group comparison and multiple comparison correction (**B,C,D,E**)

**Figure S2:**
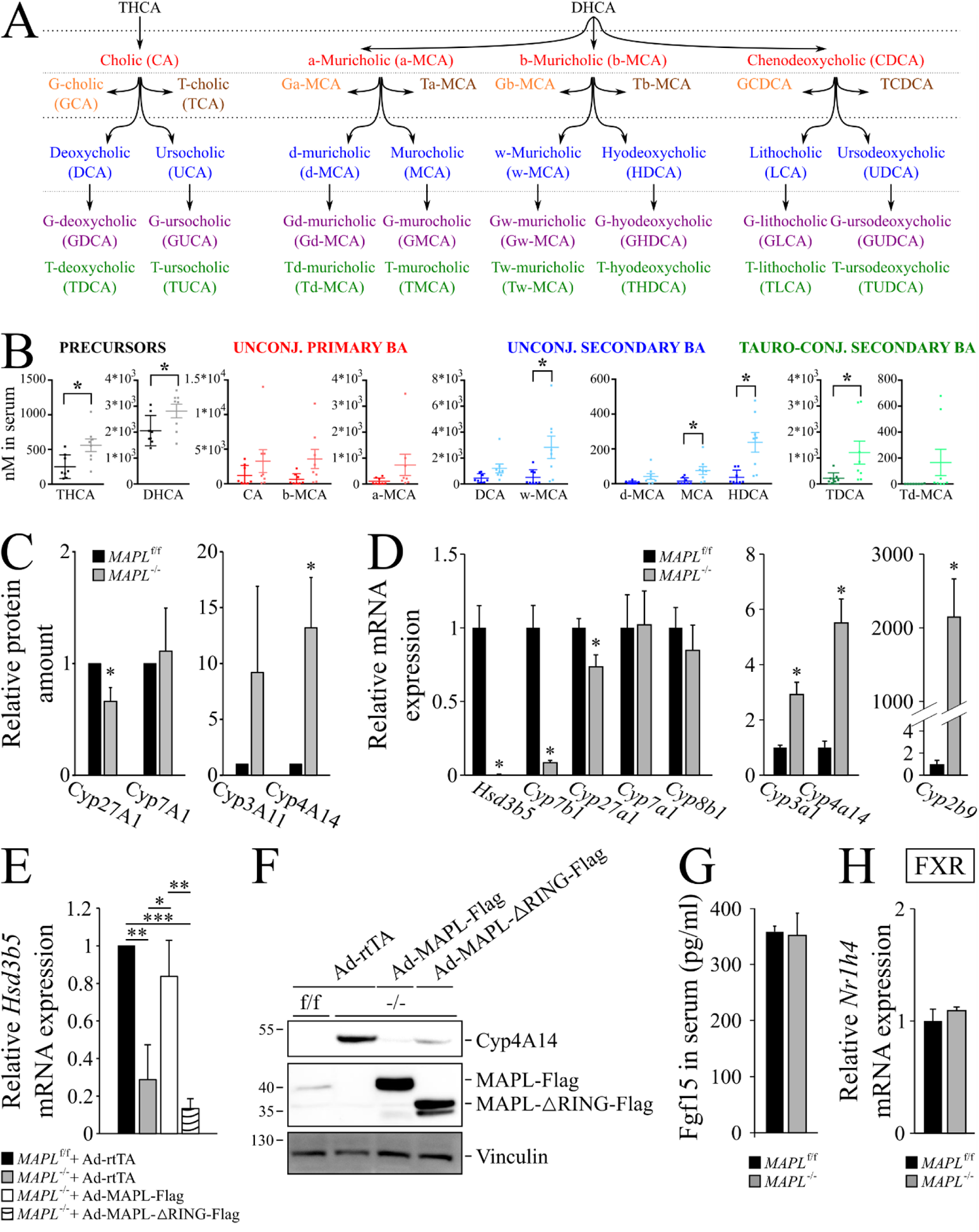
Loss of MAPL alters bile acid synthesis. **A.** A model depicting the generation of the main murine bile acids from THCA and DHCA (tri- and dihydroxycholestanoic acids) (G: glyco-; T: Tauro-). **B.** Bile acids precursors, as well as unconjugated and conjugated primary and secondary bile acids were quantified from serum (n=8 for each strain, 2-month-old males). THCA and DHCA: tri- and dihydroxycholestanoic acid; CA: cholic acid; a-MCA, b-MCA, w-MCA and d-MCA: α-, β-, γ- and δ-muricholic acid; DCA: deoxycholic acid; MCA: murocholic acid; HDCA: hyodeoxycholic acid; TDCA: taurodeoxycholic acid; td-MCA: tauro-δ-muricholic acid. **C.** Quantification of CYP27A1, Cyp7A1, Cyp3A11 and Cyp4A14 of Figure 2G western blots (n=3 for each strain). **D.** Transcriptome analysis (Illumina) was validated by qRT-PCR on livers isolated from 3 different mice for each strain (2-month-old males). **E.** qRT-PCR investigating *Hsd3b5* mRNA levels in livers of rescued mice (in triplicate, n=7 with 4 females and 3 males, n=7 with 4 females and 3 males, n=5 with 3 females and 2 males and n=6 with 3 females and 3 males, for MAPL^f/f^ + rtTA, MAPL^−/−^ + rtTA, MAPL^−/−^ + MAPL-Flag and MAPL^−/−^ + MAPL-ΔRING-Flag, respectively, 2–3-month-old). **F.** Representative western-blot of Cyp4A14 protein levels in livers isolated from rescued mice. **G.** Circulating Fgf15 was measured by ELISA (n=5 and n=6 for MAPL^f/f^ and MAPL^−/−^ respectively, 2-month-old males). **H.** qRT-PCR investigating *Nr1h4* mRNA levels (FXR) in livers of 3 different mice (2-month-old males). * *P* < 0.05 ** *P* < 0.01 *** *P* < 0.001 using T test for two group comparison and multiple comparison correction

**Figure S3:**
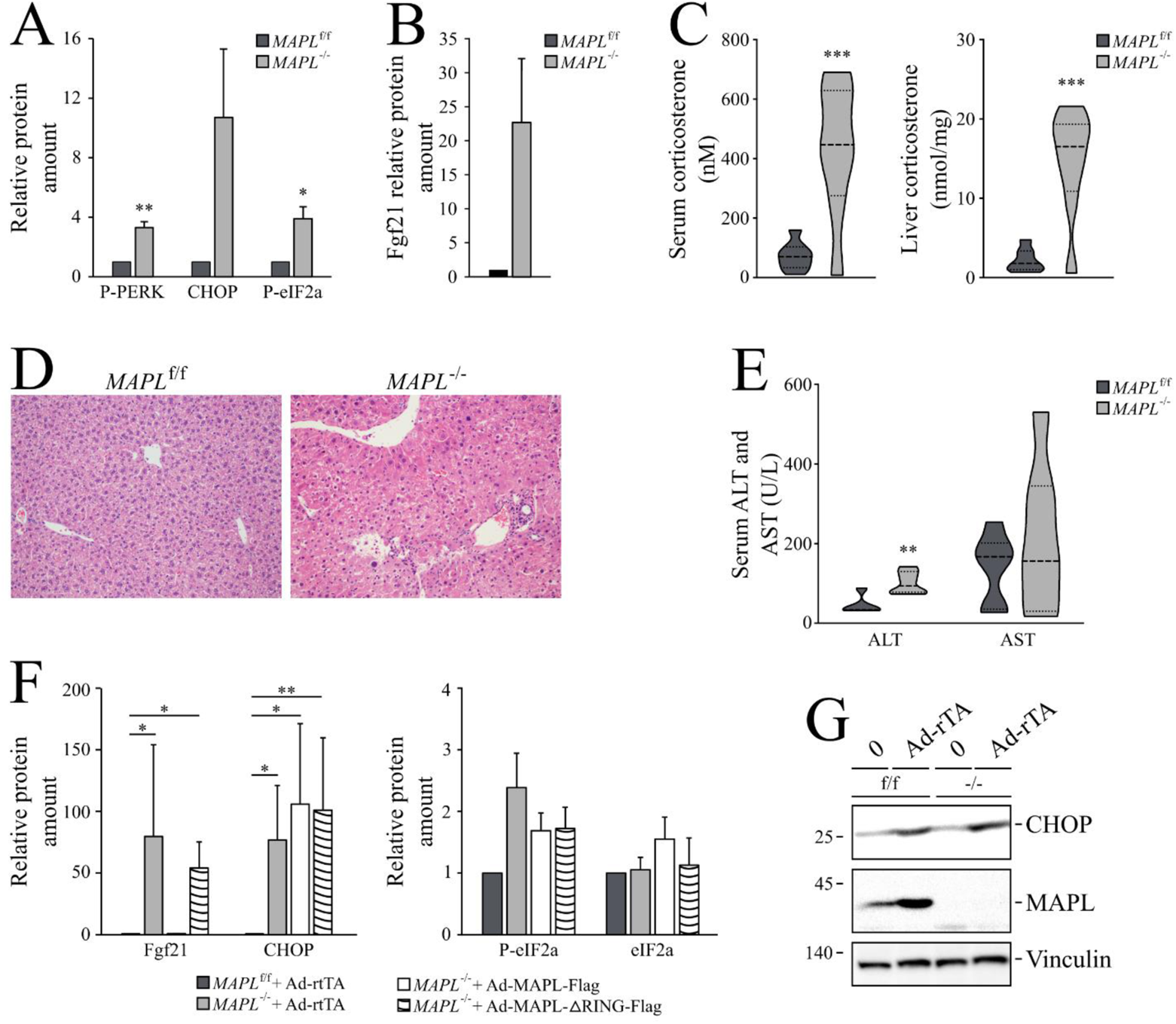
Loss of MAPL leads to hepatic ER stress, eIF2α activation and Fgf21 expression. **A.** Quantification of phopsho-PERK, CHOP and eIF2α signals presented in Figure 3A and 3B western-blots (n=3 for each strain). **B.** Quantification of Fgf21 signals presented in Figure 3D (n=3 for each strain). **C.** Circulating and liver corticosterone levels were quantified as described in materials and methods (n=8 for each strain, 2 month old males). **D.** Representative pictures of livers from *MAPL*^f/f^ and *MAPL*^−/−^ mice. Hematoxylin and eosin staining. 40X objective. **E.** Levels of alanine transaminase (ALT) and aspartate transaminase (AST), markers of liver damage, were quantified and plotted from serum collected from 6 month old females (n=5 and n=6 for *MAPL^f/f^* and *MAPL^−/−^*, respectively for ALT quantification and n=6 and n=7 for *MAPL^f/f^* and *MAPL^−/−^*, respectively for AST quantification). **F. Left panel.** Quantification of FGF21 signals presented in Figure 3E (n=5, 5, 3 and 4 for MAPL^f/f^ + rtTA, MAPL^−/−^ + rtTA, MAPL^−/−^ + MAPL-Flag and MAPL^−/−^ + MAPL-ΔRING-Flag, respectively, 2-3 month old). **Right panel**. Quantification of CHOP, P-eIF2α and eIF2α signals of Figure 3J (n=4, 4, 3 and 4 for MAPL^f/f^ + rtTA, MAPL^−/−^ + rtTA, MAPL^−/−^ + MAPL-Flag and MAPL^−/−^ + MAPL-ΔRING-Flag, respectively, 2-3 month old). **G.** Representative western-blot of CHOP protein expression of livers isolated from MAPL^f/f^ and MAPL^−/−^ animals injected or not with the empty virus rtTA. * *P* < 0.05 ** *P* < 0.01 *** *P* < 0.001 using T test for two group comparison and multiple comparison correction (**A,B,C,E**) or ANOVA for multiple group comparisons (**F**)

**Figure S4:**
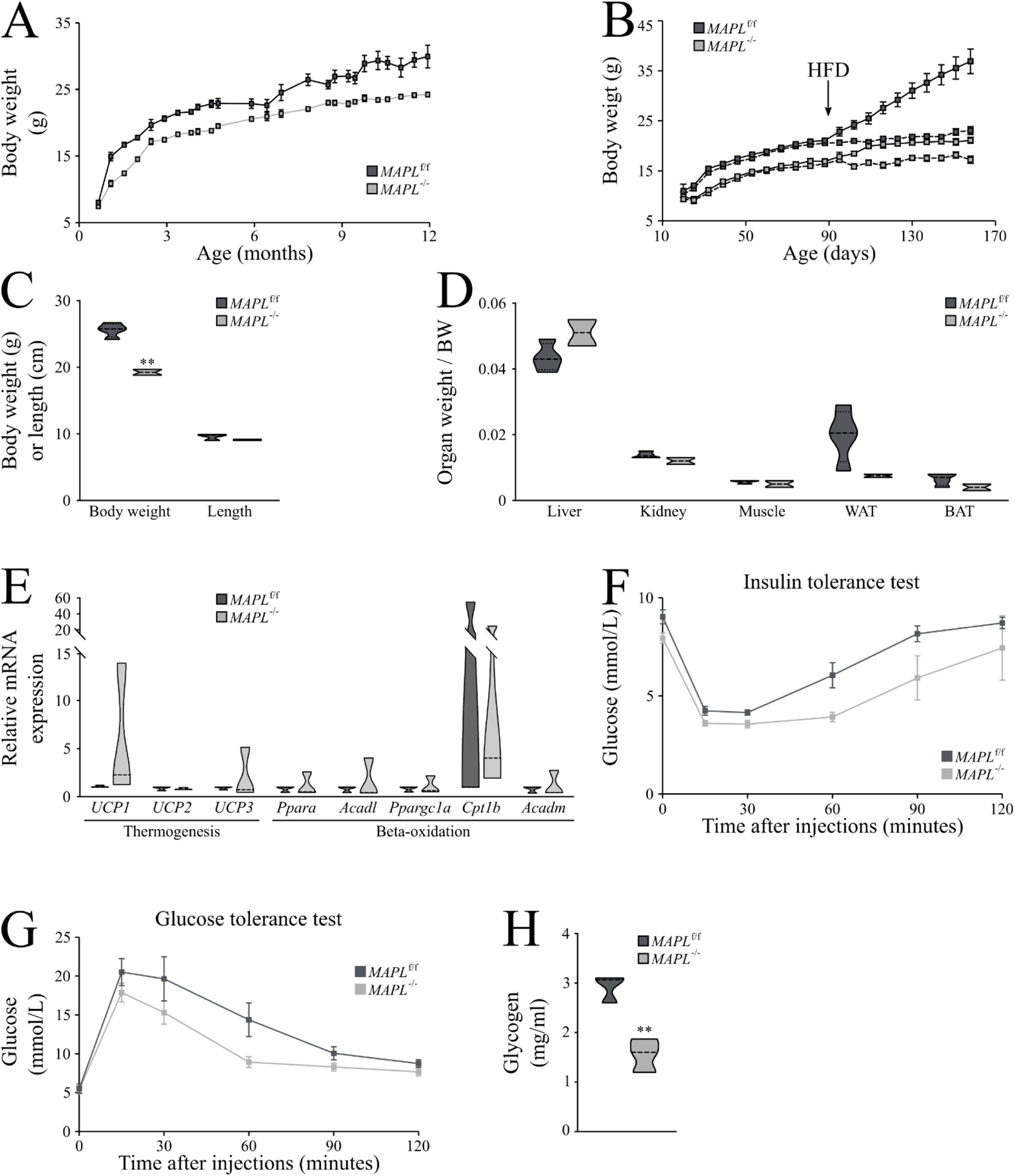
MAPL^−/−^ female mice are lean and resistant to weight gain. **A.** Female mice fed with normal chow were weighed weekly; MAPL^f/f^ (n=7) and MAPL^−/−^ (n=10). **B.** MAPL^f/f^ and MAPL^−/−^ (n=8 for each strain) females were fed with normal chow for 5 months (dotted lines), or for 3 months followed by 2 months of 60% fat chow (solid lines, diet change indicated by HFD arrow). **C.** Body weight (g, left panel) and length (cm, right panel) of 7-month old MAPL^f/f^ (n=4) and MAPL^−/−^ (n=2) female mice. **D.** Wet weight of organs including liver, kidney, gastrocnemius muscle, epididymal white fat (WAT) and interscapular brown fat (BAT) isolated from 7-month-old MAPL^f/f^ (n=4) or MAPL^−/−^ (n=2) female mice. **E.** White adipose tissue gene expression performed by qRT-PCR. N=3, in triplicate, 2-month-old MAPL^fl/fl^ and MAPL^−/−^ male mice. **F.** Insulin tolerance test in MAPL^f/f^ (n=7), MAPL^−/−^ (n=11) female mice. Insulin (0.5 U/kg) was injected intraperitoneally after a 2 h fast and blood glucose was measured at indicated times. **G.** Glucose tolerance test in MAPL^f/f^ (n=7), MAPL^−/−^ (n=11) female mice. Glucose (2 g/kg) was injected intraperitoneally after an overnight fast and blood glucose was measured at indicated times. **H.** Liver glycogen measured enzymatically in female *MAPL*^f/f^ (n=7), *MAPL*^−/−^ (n=11) mice. * *P* < 0.05 ** *P* < 0.01 *** *P* < 0.001 using T test for two group comparison and multiple comparison correction or ANOVA for multiple group comparisons

**Figure S5:**
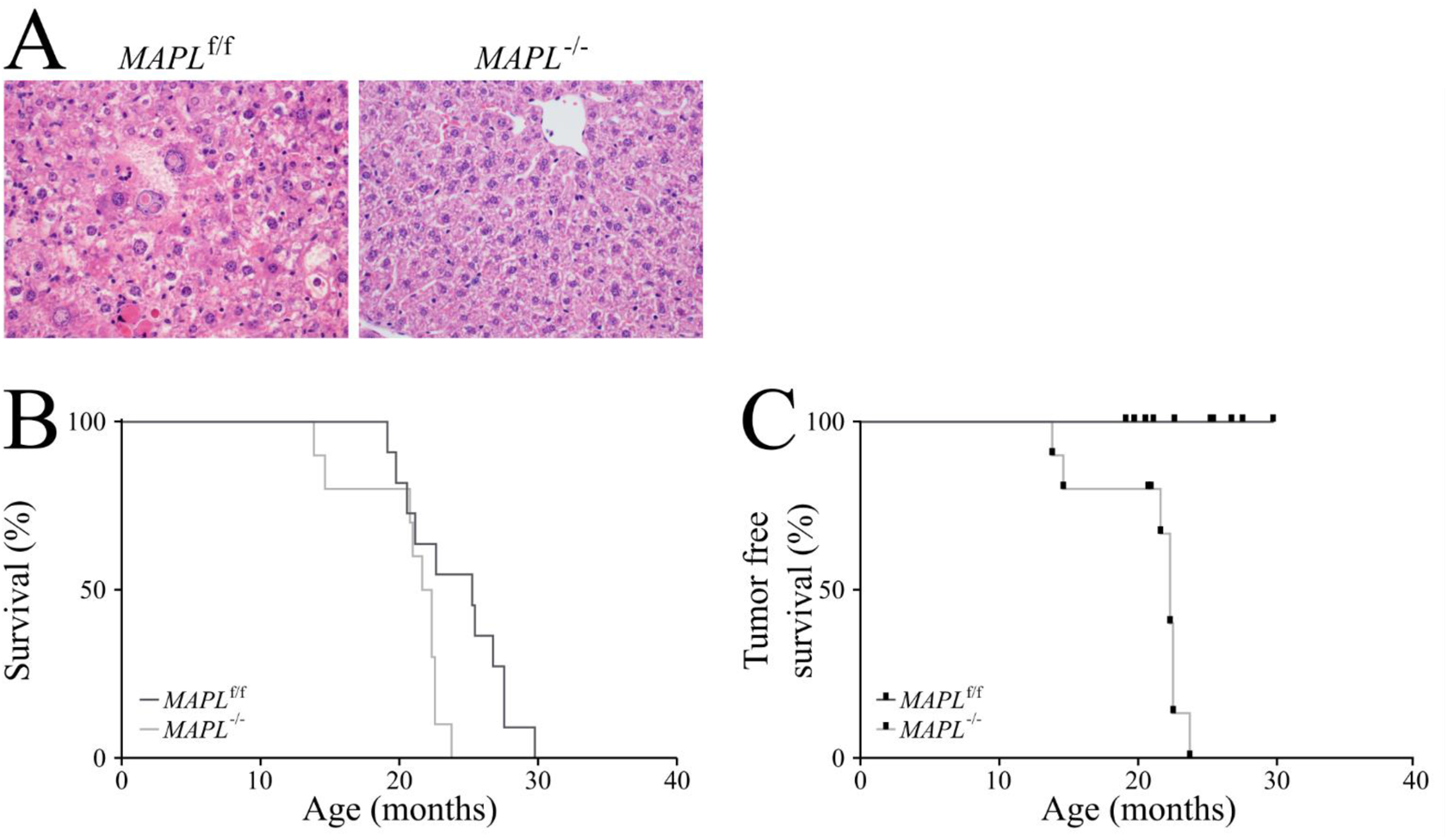
Spontaneous development of hepatocellular carcinoma in female MAPL deficient mice. A. Hematoxilin eosin staining of female MAPL^f/f^ and MAPL^−/−^ livers. B. Survival curve of MAPL^f/f^ (N=11) and MAPL^−/−^ (N=11) female mice. C. Cancer free survival curve of MAPL^f/f^ (N=11) and MAPL^−/−^ (N=11) female mice.

**Supplementary Table 1: BioID analysis of MAPL interacting proteins.**

MAPL-BirA*Flag BioID results. Data are presented as spectral counts detected for each prey protein, as indicated. Two technical replicates were performed on each of two unique biological replicates (for a total of four MS analyses). Selection of preys was based on ProteinProphet confidence score *p*≥0.85 (FDR <1%) and SAINT Express score ≥0.9. For control runs, only the highest four spectral counts (out of 16 runs) are shown.

**Supplementary Table 2: Quantification of bile acid species from liver and serum.**

Precursors, unconjugated and conjugated primary and secondary bile acids were quantified as described in materials and methods on liver and serum from 2-month-old males (n=8 for each strain).

**Supplementary Table 3: Transcriptome analysis from liver.**

Table presenting data from the Illumina analysis performed on 5-month-old males (n=3, for each strain, in triplicate) as described in material and methods, summarizing genes with variations higher than 2-fold (with a p value lower than 0.05) implicated in different metabolic pathways.

## Notes

The authors have no conflict of interests to declare

